# A cafeteria diet blunts effects of exercise on adult hippocampal neurogenesis but not neurogenesis-dependent behaviours in adult male rats

**DOI:** 10.1101/2024.07.16.603714

**Authors:** Minke H.C. Nota, Sebastian Dohm-Hansen, Sarah Nicolas, Erin P. Harris, Tara Foley, Yvonne M. Nolan, Olivia F. O’Leary

## Abstract

Animal studies have shown that a cafeteria (CAF) diet (high in saturated fat and sugar), is associated with memory impairments and increased anxiety, while exercise can enhance antidepressant-like effects and cognitive function. The mechanisms underlying the effects of a CAF diet, exercise, or their convergence on memory, mood and anxiety are not fully understood, but alterations in adult hippocampal neurogenesis (AHN), gut microbial metabolites, or plasma metabolic hormones may play a role. Therefore, this study investigated whether a 7.5-week voluntary running exercise intervention in young adult male rats could alter the effects of a concurrent CAF diet on depression-like, anxiety-like and cognitive behaviours and AHN, and determined associated changes in metabolic hormones and gut microbial metabolites. We found that exercise produced a mild anxiolytic effect, regardless of diet, and increased PYY, a hormone previously shown to reduce anxiety-like behaviour. CAF diet induced differential abundance of caecal metabolites, and exercise attenuated CAF diet-induced decreases in certain metabolites implicated in cognitive function or depression-like behaviour. Although exercise exerted antidepressant-like effects in the FST, induced subtle improvements in spatial learning strategy, and increased plasma metabolic hormones previously implicated in depression-like behaviour in CAF diet-fed animals, CAF diet blunted exercise-induced increases in plasma GLP-1 and AHN, suggesting that exercise should be accompanied by a healthy diet to increase AHN. Together, these findings highlight the importance of exercise and healthy diet for hippocampal health and provide insight into potential metabolite and hormone-mediated mechanisms underlying the effects of CAF diet and exercise on brain and behaviour.

**Key points:**

- Diets high in saturated fat and sugar are associated with memory impairments and increased anxiety while exercise can exert antidepressant-like effects and enhance cognitive function, but the biological underpinnings of these effects and whether exercise can negate effects of such diets remain to be elucidated.
- We found that running exercise modestly reduced anxiety in rats fed either a healthy or a cafeteria-style diet and increased a hormone (peptide YY) previously shown to decrease anxiety.
- Running exercise exerted antidepressant-like effects in cafeteria diet-fed rats and attenuated cafeteria diet-induced decreases in gut metabolites previously implicated in cognition or depression-like behaviour.
- Cafeteria diet blunted exercise-induced production of new neurons in the hippocampus, a brain region important in mood and memory.
- These data highlight the importance of combining exercise with a healthy diet for hippocampal health, while identifying potential targets for intervention or dietary supplementation to prevent a cafeteria diet blunting beneficial effects of exercise

## 1. Introduction

There is a global increase in the availability of highly processed, energy-dense foods (da Costa *et al*., 2022), as well as sedentary lifestyles (Park *et al*., 2020). A poor diet, such as a Western-style diet high in saturated fats and sugar, and sedentary behaviour can lead to obesity and subsequent metabolic alterations (Singla *et al*., 2010; Rakhra *et al*., 2020; Silveira *et al*., 2022) including alterations in circulating metabolic hormones. These negative lifestyle factors are known to increase the risk of depression and anxiety (Hryhorczuk *et al*., 2013), which significantly impacts quality of life and in severe cases can lead to suicide (World Health Organisation, 2021). In addition, metabolic alterations as a result of poor diet and sedentary behaviour have been associated with cognitive impairment (Duarte, 2015; Kordestani-Moghadam *et al*., 2020). Cognitive impairment is highly correlated with depression and significantly contributes to functional debilitation (Perini *et al*., 2019). There is accumulating evidence of bidirectional communication between gut resident microbiota and the brain, known as the microbiota-gut-brain axis (MGBA) (Cryan *et al*., 2019). Indeed, neuropsychiatric disorders such as depression and cognitive disorders such as Alzheimer’s disease are associated with altered gut microbiota composition (Donoso *et al*., 2022; Grabrucker *et al*., 2023) and there is evidence that lifestyle factors including diet and exercise can influence the MGBA (Nota *et al*., 2023). Given that the gut is at the interface between diet and host and can modulate brain function via the MGBA, and that diet and exercise influence the MGBA (Guzzetta *et al*., 2022; Nota *et al*., 2023), alterations in gut microbial metabolites may be a plausible mechanism through which Western-style diets and exercise can influence brain and behaviour.

Reports from animal studies have shown negative effects of diets high in fat and/or sugar on depression-like behaviour, anxiety-like behaviour, and cognitive behaviours such as working and object recognition memory, and spatial learning and memory. Obesity induced by a 2-month high fat diet (HFD) increased anxiety-like behaviour and decreased reward-seeking behaviour in adult male mice (Li *et al*., 2022). The impact of a cafeteria (CAF) diet, which is high in saturated fat and sugar and mimics human Western-style diets, on such behaviours has been less investigated. However, it has been reported that a CAF diet increased insulin concentrations and decreased spatial and recognition memory in adult male rats after 20 weeks (Lewis *et al*., 2019), and impaired spatial memory and increased anxiety-like behaviour in adolescent male rats after 12 weeks (Ferreira *et al*., 2018). Conversely, research in rodents has demonstrated that exercise has beneficial effects on anxiety-like and depression-like behaviour, pattern separation, and spatial learning and memory (Caruso *et al*., 2024). For example, 3-4 weeks of voluntary running exercise decreased learned helplessness and anxiety-like and depression-like behaviour in adult male mice (Duman *et al*., 2008) and mice with access to voluntary running exercise for 4 weeks showed improvements in spatial learning and memory (Liu *et al*., 2009). Furthermore, 7 weeks of voluntary wheel running enhanced contextual and cued fear conditioning (O’Leary *et al*., 2019*a*), and 11-12 weeks of voluntary running exercise improved reversal learning in a touchscreen location discrimination task in both adolescent and adult male rats (O’Leary *et al*., 2019*b*). However, whether exercise can mitigate CAF diet-induced effects on depression-like, anxiety-like and cognitive behaviours and circulating metabolic hormones has not yet been determined.

Dysfunction of the hippocampus is a hallmark feature of depression, anxiety, and cognitive impairment (de Asis *et al*., 2001; de Rover *et al*., 2011; Tural *et al*., 2018). Adult hippocampal neurogenesis (AHN), the birth of new neurons in the dentate gyrus of the hippocampus, is known to play a role in anxiety-like behaviour (Revest *et al*., 2009) and cognitive functions such as pattern separation (Clelland *et al*., 2009; Bekinschtein *et al*., 2014) and spatial memory (Shimazu *et al*., 2006; Zhang *et al*., 2008). Furthermore, AHN is required for some of the behavioural and endocrine effects of certain antidepressants (Santarelli *et al*., 2003; Surget *et al*., 2011). Critically, AHN is sensitive to external factors including diet and exercise (Hueston *et al*., 2017). A 12-week CAF diet fed from adolescence decreased AHN in male rats (Ferreira *et al*., 2018; Liśkiewicz *et al*., 2021), but the effects of adult-initiated CAF diet on AHN are not yet well-characterised. On the other hand, beneficial effects of exercise on AHN have been reported (van Praag *et al*., 1999). A 7-week voluntary running exercise intervention has been demonstrated to increase AHN in adult male rats (Nokia *et al*., 2016). In addition, in male rats, both adult- and adolescent-initiated exercise (11 and 12 weeks, respectively) have been shown to increase neurite complexity of maturing hippocampal neurons (O’Leary *et al*., 2019*b*). Importantly, metabolic hormones, such as leptin (Calió *et al*., 2021), ghrelin (Moon *et al*., 2009), insulin (Bayram *et al*., 2022), glucagon-like peptide 1 (GLP-1) (Isacson *et al*., 2011), and fibroblast growth factor 21 (FGF-21) (Wang *et al*., 2018) can be altered as a result of diet or exercise and can also impact AHN, suggesting possible mechanisms through which diet and exercise might regulate AHN and AHN-associated behaviours.

The MGBA has also been reported to influence AHN and related behaviours (Ogbonnaya *et al*., 2015; Cruz-Pereira *et al*., 2020; Cavallucci *et al*., 2020), through various mechanisms including production of microbial metabolites (Morais *et al*., 2021; Guzzetta *et al*., 2022). Essential for host homeostasis, microbial metabolites are small compounds produced by intestinal microorganisms, and include amongst others short-chain fatty acids (SCFAs), (essential) amino acids, and neurotransmitters (Krautkramer *et al*., 2021). Dietary components such as fibre and lipids play an important role in the modulation of gut microbial metabolism (Nishida *et al*., 2022) and indeed, a recent study indicated that switching from standard chow to a HFD altered caecal metabolite composition in adult male rats (Fouesnard *et al*., 2021). However, it is unknown whether a CAF diet induces similar changes. Conversely, exercise has been indicated to produce numerous beneficial effect on gut health, for example by increasing gut microbiota diversity (Mitchell *et al*., 2019; Xu *et al*., 2022) and SCFA production (Barton *et al*., 2018), and it has been demonstrated that 3-6 months of treadmill exercise attenuated HFD-induced as well as toxin exposure-induced gut microbiota dysbiosis in adolescent male mice (Wang *et al*., 2022; Fu *et al*., 2022). Notably, a voluntary exercise-induced increase in AHN was attenuated when gut microbiota were depleted using chronic administration of antibiotics (Nicolas *et al*., 2024), suggesting a role for the gut microbiota in the pro-neurogenic effects of exercise.

It remains unclear if exercise can attenuate any negative effects of a CAF diet on AHN and hippocampus-associated anxiety-like, depression-like and cognitive behaviours. Furthermore, the interactions between a CAF diet and exercise on metabolic hormones and microbial metabolite production are not well-known. Therefore, this study investigated whether a 7.5-week voluntary running exercise intervention in young adult male rats altered the effects of a concurrent CAF diet on depression-like and anxiety-like behaviours, pattern separation, recognition memory and spatial learning, as well as AHN. In addition, whether alterations in behaviour and AHN were associated with changes in metabolic hormones and gut microbial metabolites was investigated.

## 2. Methods

### 2.1 Animals

Male Sprague-Dawley rats were obtained from Envigo Laboratories (United Kingdom) at approximately 7 weeks old (225-250g) and housed in groups of four in standard conditions (22 ± 1°C, 50% relative humidity) on a 12-hour light-dark cycle (lights on at 7:00am), with *ad libitum* access to food and water. At the start of the experiment, at approximately 9 weeks old, animals were pair-housed for the duration of the study. All animal procedures were performed under licenses issued by the Health Products Regulatory Authority (AE19130/P123, HPRA, Ireland), in accordance with the European Communities Council Directive (2010/63/EU) and approved by the Animal Experimentation Ethics Committee of University College Cork (2019/025).

### 2.2 Experimental design

Animals were pair-housed and randomly divided into four experimental groups; sedentary animals with access to standard chow (CTRL-SED, n=12), sedentary animals with access to cafeteria diet (CAF-SED, n=12), animals with voluntary access to running wheels and standard chow (CTRL-EX, n=12), and animals with voluntary access to running wheels and cafeteria diet (CAF-EX, n=12, Figure 1A). Standard chow (Envigo, UK) consisted of 6.2% fat (of which 0% saturated fat), 44.2% carbohydrates (of which 0% sugar), 18.6% protein, and 3.5% fibre (Supplementary Table 1). The cafeteria diet consisted of several different food items high in fat and/or sugar, with two high-fat and two high-sugar items given each day in rotation in addition to standard chow, for the duration of the experiment (7.5 weeks), as previously described by Nicolas *et al*. (2022) (Supplementary Table 1). All food was provided in excess, *ad libitum*. Exercising animals had continuous access to a running wheel (Techniplast, UK) for the duration of the experiment. Running distance (km) was recorded in 24h increments. Weight gain was calculated as 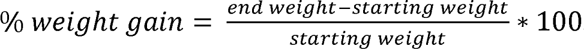. There was no significant difference in weight between groups at the start of the experiment.

**Figure 1:**
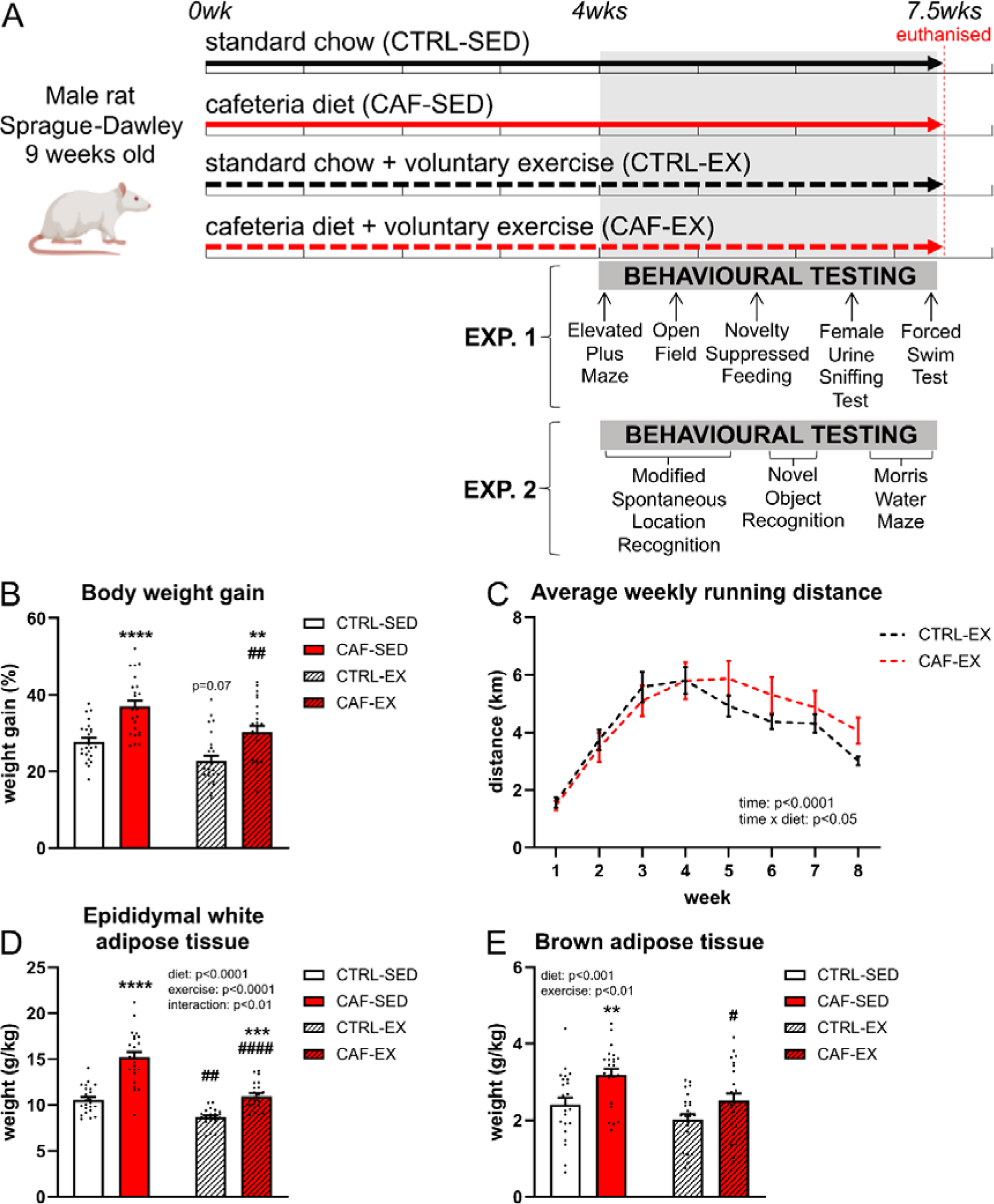
**(A)** Experimental design. **(B)** Total body weight gain (%) compared to starting weight (n=24, exp. 1&2). **(C)** Average weekly running distance (km, n=12, exp. 1&2). **(D)** Effects of a CAF diet and exercise on body weight-adjusted (g/kg) epidydimal white adipose tissue (eWAT) weight (n=23-24, exp. 1&2). **(E)** Effects of a CAF diet and exercise on body weight-adjusted (g/kg) brown adipose tissue (BAT) weight (n=23-24, exp. 1&2). All data are expressed as mean ±SEM, **p<0.01, ***p<0.001 & ****p<0.0001 vs. corresponding treatment group with access to standard chow; p=0.07, #p<0.05, ##p<0.01 & ####p<0.0001 vs. corresponding sedentary treatment group.

Four weeks after the start of the interventions, anxiety-like behaviour was assessed in the Elevated Plus Maze and Novelty Suppressed Feeding tests, anhedonia in the Female Urine Sniffing test and antidepressant-like behaviour in the Forced Swim test (Figure 1A). In a separate cohort of rats, which were randomly allocated to the same experimental groups (CTRL-SED, n=12; CAF-SED, n=12; CTRL-EX, n=12; CAF-EX, n=12), pattern separation, recognition memory, and spatial learning and memory were assessed in the Modified Spontaneous Location Recognition, Novel Object Recognition, and Morris Water Maze test, respectively (Figure 1A). These behavioural tests were likewise conducted four weeks after the start of the interventions.

Half of the animals in each group (n=6) were transcardially perfused 1-3 hours after cessation of the diet and exercise exposures for subsequent immunohistochemical assessment of the survival of newly born neurons, as well as the number of immature hippocampal neurons as a measure of AHN. The other half of the animals in each group (n=6) of each of the two cohorts were euthanised by rapid decapitation 1-3 hours after cessation of the diet and exercise exposures for collection of fresh tissue (adipose tissue to determine adiposity, trunk blood for metabolic hormone analysis, and caecum for metabolomic analysis).

### 2.3 Behavioural testing

#### 2.3.1 Elevated Plus Maze

Anxiety-like behaviour was assessed in the Elevated Plus Maze (EPM) according to Harris et al., 2022b. Animals were habituated to the testing room 1h prior to commencing the test. The dimly lit (red light, ±5lux) maze consisted of two opposed open (50cm length x 10cm width x 25cm height) and two opposed closed arms (50cm length x 10cm width x 40cm height) mounted at a 90° angle, all facing a central platform (10cm x 10cm), elevated 50cm above the floor. Each rat was placed in the centre of the EPM facing an open arm and left to explore freely. Their behaviour was monitored and video tracked for 5min, after which the animal was returned to its home cage. The maze was cleaned using 70% ethanol between each animal to eliminate olfactory cues. Time spent in each portion of the maze (open arms, closed arms, centre), and entries in the open and closed arms, were scored manually and blinded to experimental groups. Arm entries were defined as all four paws of the animal being across the borders of the arm. Data are presented as the percentage of total test time in open arms and closed arms.

#### 2.3.2 Novelty Suppressed Feeding

The Novelty Suppressed Feeding (NSF) test was used to measure anxiety-like behaviour (Harris *et al*., 2022*b*). The day before the test (6p.m.), all food was removed from the home cage. Animals were food-deprived for no more than 16h. The day of the test, animals were habituated to the experimental room for 1h. Rats were then placed in a brightly lit (±1000lux) open field arena with bedding inside. A food pellet was placed on a white plastic base in the centre of the arena. Latency in time (s) to begin eating was recorded during 10min. As soon as the rat was observed to eat, or the 10min time limit was reached, the rat was removed from the open field and returned to the home cage where it had access to pre-weighed standard chow. After 30min, chow was removed and weighed to determine the amount consumed by the rat, which was adjusted to body weight (g/kg). The arena and food platform were cleaned using 70% ethanol between each animal to eliminate olfactory cues.

#### 2.3.3 Female Urine Sniffing Test

Anhedonia was measured using the Female Urine Sniffing Test (FUST) according to Harris et al., 2022b. Urine was collected from adult female Sprague-Dawley rats in oestrous. The cycle was determined by collecting vaginal secretion with a plastic transfer pipette (tip diameter <1mm) onto a glass slide. The secretion was observed under a light microscope to determine the phase of oestrous cycle.

Experimental male animals were habituated to the testing room for 1h. During the last 45min of habituation, a clean, dry cotton bud was placed inside each cage. The cotton bud was removed and each animal was exposed to a new cotton bud with dH_2_O for 3min, after which it was removed. Following a 45min interval, the animal was then exposed to another cotton bud with female oestrus urine for 3min. Each exposure was recorded by video camera, and time spent sniffing the cotton buds was scored manually and blinded to experimental groups. Preference to sniff urine compared to water was calculated as 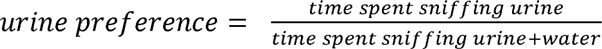.

#### 2.3.4 Forced Swim Test

The modified Forced Swim Test (FST) was used to measure depression-like behaviour (Porsolt *et al*., 1978). In the pre-swim test, rats were individually exposed to a water tank for 15min (21cm diameter, filled to 30cm with 23-25°C water). Twenty-four hours later, rats were placed back in the water tank for 5min. Behaviour during the 5min test was recorded by a video camera. Following the test, the rat was removed from the tank, gently towel-dried and returned to its home cage. Active (climbing and swimming), and passive (immobility) behaviours during the 5min swim were scored manually and blinded to experimental groups in 5s time bins as previously described (Slattery & Cryan, 2012).

#### 2.3.5 Open Field

General locomotor activity was assessed in the Open Field test. Animals were habituated to the testing room for 1h prior to commencing the test. Each rat was placed in the centre of a brightly lit (±1000lux) arena (90cm diameter) and allowed to explore for 10min. Animals were then removed and placed back into their home cage. Distance moved (m) and time in the centre (s) were analysed using Noldus EthoVision XT 11.5 tracking software. The diameter of the centre area was set at 45cm according to (Kozareva *et al*., 2019). The arena was cleaned using 70% ethanol between each animal exposure to eliminate olfactory cues.

#### 2.3.6 Modified Spontaneous Location Recognition Test

Pattern separation, which is associated with AHN and refers to the ability to discriminate between highly similar memories, was assessed in the Modified Spontaneous Location Recognition (MSLR) test according to Kozareva et al., 2019. Animals were habituated to a dimly lit circular arena (∼20lux, 90cm diameter) with bedding for 10min per day for five consecutive days prior to testing. Proximal cues were placed in the testing room for spatial navigation. Following the last day of habituation, during the acquisition phase, animals were presented with three identical objects (33cl glass beer bottles, or soda cans) and allowed to explore them for 10min, once for the large separation (LS) test and once for the small separation (SS). For the LS paradigm, all three objects were separated by 120° (Figure 2.2B). For the SS paradigm, two of the objects (A1 and A2) were separated by 50° with the third object (A3) at an equal distance between them (Figure 2.2C). Twenty-four hours after acquisition, animals were presented with two of the objects used in the acquisition phase and allowed to explore them for 5min. The familiar (A4) was placed in the same location as A3, while the novel (A5) was placed directly between the locations of A1 and A2 (Figure 2.2B-C). Separation order, object type, and object location were randomised across tests. Behaviour was recorded and videos were analysed with blinding to experimental groups to determine exploration time with novel and familiar objects. The discrimination ratio (DR) was calculated as 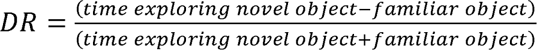. Objects were cleaned using 70% ethanol between each animal to eliminate olfactory cues.

**Figure 2:**
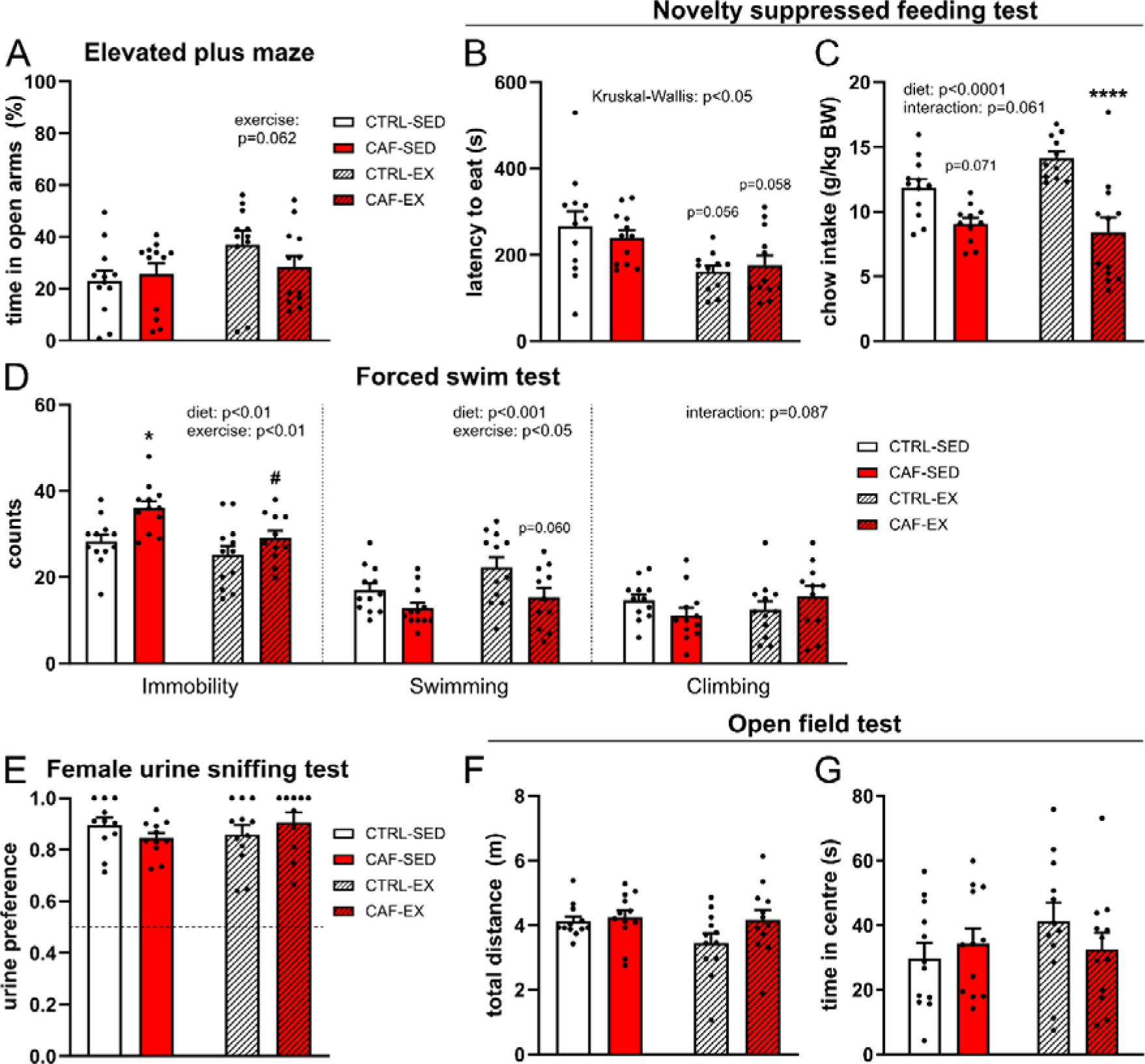
**(A)** Percentage of total time spent in the open arms of the elevated plus maze (EPM, n=11-12). **(B)** Latency (s) to begin eating standard chow pellet on the centre platform of the novelty suppressed feeding (NSF) test arena (n=11-12). **(C)** Standard chow consumption (grams) during the 30min post-testing phase (n=11-12), adjusted for body weight. **(D)** Immobility, swimming, and climbing scores in the forced swim test (FST, n=11-12). **(E)** Urine preference as time spent sniffing urine/total sniffing time in the female urine sniffing test (FST (n=10-12). **(F)** Total distance travelled (m) in the open field test (OFT, n=12). **(G)** Total time (s) spent in the centre area of the OFT arena (n=12). All data are expressed as mean ±SEM, p=0.071, *p<0.05 & ****p<0.0001 vs. corresponding treatment group with access to standard chow; p=0.060, p=0.058, p=0.056 & #p<0.05 vs. corresponding sedentary treatment group.

#### 2.3.7 Novel Object Recognition

Recognition memory was measured in the Novel Object Recognition test (NOR). Animals were habituated to a dimly lit circular arena devoid of bedding (∼20lux, 90cm diameter) for 10min. The next day, two identical objects (ceramic mug, or 250ml graduated borosilicate bottle) were placed in the arena and the animals were allowed to explore them for 10min. Twenty-four hours after acquisition, one of the objects was replaced with a novel object (the object not used during acquisition), and animals were allowed to explore for 5min (test, Figure 2.2J). Behaviour was recorded and videos were analysed with blinding to experimental groups to determine exploration time with novel and familiar objects in the 5min test. The DR was calculated as 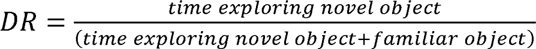. The arena and objects were cleaned using 70% ethanol between each animal to eliminate olfactory cues.

#### 2.3.8 Morris Water Maze

Hippocampal-dependent spatial learning and memory were assessed in the Morris Water Maze (MWM, Figure 2.4A) as previously described (Hoban *et al*., 2016). Prior to testing, animals were habituated to the testing room for 1h. A black, circular pool (180cm diameter) was filled with water (22 ±1°C). A hidden, transparent platform was placed 1-2cm below the water surface at a fixed location in the north-west quadrant 15cm from the pool wall. External cues were placed in the testing room for spatial navigation. During training (acquisition days 1-4), animals were placed in the pool at one of four release points (NE, E, S, and SW), with each point being used only once per day. Animals underwent one block of four training trials each day for four consecutive days, with each trial lasting a maximum of 120s. If the animal found the platform within that time, they were allowed to stay on the platform for 10s before continuing to the next trial. However, if the animal failed to reach the platform within the maximum trial time, they were guided to the platform and remained there for 30s before continuing to the next trial. All trials were recorded, and videos were analysed using Noldus EthoVision XT 11.5 tracking software. Learning performance during training was characterised by average escape latency to locate the platform, average swimming velocity, and the animal’s mean distance from the platform (whereby smaller distance indicates more movement close to the platform) each day. Spatially imprecise hippocampus-independent (thigmotaxis, scanning, and circling), and spatially precise hippocampus-dependent (focal search, rotating, and direct swim) search strategies during training were assessed according to Higaki et al., 2018 and Gil-Mohapel et al., 2013. The swimming path in each trial was traced using ANY-maze video tracking software system and manually assigned a cognitive score between 0-5, with 0=thigmotaxis; 1=scanning; 2=circling; 3=focal search; 4=rotating; and 5=direct swim (Figure 2.5A), blinded to experimental groups. The average daily cognitive score was calculated as the average of all four trials per day.

On the fifth day (probe trial), the platform was removed and animals were released in the pool from the SE point. Animals were allowed to explore the pool for 60s. Latency to enter and time spent in the platform quadrant, and average velocity were analysed using Noldus EthoVision XT 11.5 tracking software to assess spatial memory.

### 2.4 Blood and tissue collection

One day after the last behavioural test, 1-3 hours after cessation of the diet and exercise exposures, half of the animals in each group (n=6) were euthanised by rapid decapitation. Trunk blood was collected in 3ml EDTA-coated tubes (Vacuette®, Greiner Bio-One). EDTA-collected blood samples were centrifuged for 10min at 3220xg at 4°C to collect plasma, which was stored at −80°C until used for measurement of metabolic hormones. Animals were weighed prior to euthanasia, and epidydimal white adipose tissue (eWAT) and brown adipose tissue (BAT) were dissected to determine adiposity (grams eWAT or BAT per kg body weight). Whole caecum (n=12) was collected and weighed to determine caecum weight per kg body weight. Caecum content (n=10) was snap frozen using dry ice and stored at −80°C until preparation for metabolomic analysis.

The other half of the animals in each group (n=6) were weighed and underwent transcardial perfusion 1-3 hours after cessation of the diet and exercise exposures. Sodium pentobarbital (90mg/kg) was administered by intraperitoneal injection as anaesthetic overdose. Sufficient depth of anaesthesia was identified by loss of toe pinch reflex. Animals were placed in a dorsal recumbent position with their paws pinned. The abdomen of the animals was opened, and a catheter was inserted directly into the heart left ventricle. A cut was made in the right side of the heart to allow efflux of blood and injected fluids. Ice-cold phosphate-buffered saline (PBS) was perfused using a pump (35-40mL/min) until efflux ran clear, followed by 4% paraformaldehyde (PFA) in PBS until full body stiffness was reached. During the PBS step, epidydimal white adipose tissue was dissected and weighed to determine adiposity. Whole brains were removed and collected in 4% PFA in PBS for post-fixation. After 24h, brains were transferred to 30% sucrose in PBS until fully sunken. Brains were then snap frozen in isopentane using liquid nitrogen and stored at −80°C until ready for sectioning. Fixed brains were sectioned coronally at 40μm using a Leica CM1950 cryostat, collected free-floating in a series of twelve in cryoprotectant (25% 0.1M PBS, 30% ethylene glycol, 25% glycerol, 20% dH_2_O) and stored at −20°C until immunohistochemical staining.

### 2.5 Immunohistochemistry

For immunohistochemical staining of doublecortin (DCX, n=6 per group), a marker of immature neurons, sections were washed in 0.1M PBS (3×5min) and placed in blocking solution (10% donkey serum in 0.5% Triton X-100 in PBS (PBS-T)) for 2h at room temperature (RT). Sections were then incubated in primary antibody (rabbit anti-DCX; Abcam, AB18723, 1:5000) for 48h at 4°C. All antibodies were diluted in 5% donkey serum in 0.5% PBS-T. Sections were washed in 0.5% Tween in PBS (3×20min). All subsequent steps were carried out protected from light. Sections were incubated in secondary antibody (Alexa Fluor 488-conjugated donkey anti-rabbit, Invitrogen, AB21206, 1:500) for 2h at RT, and washed again in 0.5% Tween in PBS. Sections were incubated in DAPI (5mg/ml, 1:50000 in PBS) for 3min and washed in PBS, then mounted onto Superfrost Plus slides and cover-slipped using Dako fluorescent mounting media. Slides were stored in the dark at 4°C until imaging.

### 2.6 Microscopy and image analysis

The dentate gyrus was imaged using an Olympus BX53 Upright Research Microscope at 20x magnification for DCX. The number of DCX^+^ cells in the dentate gyrus were counted in every twelfth section using ImageJ and blinded to experimental groups. Dentate gyrus area (mm^2^) was measured using ImageJ on 10x magnification images of DAPI, respectively, and cell count results were expressed as cells/mm^2^. The dorsal hippocampus (dHi) was defined as AP: Bregma −1.8 to −5.2, and ventral hippocampus (vHi) as AP: −5.2 to −6.7 (Brummelte & Galea, 2010; Ramos Costa *et al*., 2019; Harris *et al*., 2022*a*). Sections at AP: −5.2 were only considered as vHI if the ventral part of the dentate gyrus was clearly present at the bottom of the section (Brummelte & Galea, 2010). Note: in a coronally sectioned rat brain, rostral sections typically only contain dHi, whereas caudal sections containing vHi may also contain portions of dHi and intermediate hippocampus (Tanti & Belzung, 2013; O’Leary & Cryan, 2014). For analysis, 3 dorsal and 3 ventral sections were used, taking care to select sections of similar AP coordinates from Bregma for all animals where possible.

### 2.7 Plasma metabolic hormone measurements

Concentrations of plasma hormones were measured in duplicate by ELISA. A U-PLEX Custom Metabolic Group 1 assay (K153ACL-1, MesoScale Discovery) was used to measure C-Peptide (220 - 125,000 pg/mL), fibroblast growth factor (FGF) 21 (2.8 - 8,230 pg/mL), total ghrelin (1.7 - 2,710 pg/mL), total glucagon-like peptide (GLP) 1 (0.59 – 576 pM), glucagon (0.13 – 156 pM), insulin (3.0 - 5,500 μIU/mL), leptin (11 - 50,000 pg/mL), and total peptide YY (PYY, 1.1 - 4,000 pg/mL) according to manufacturer guidelines.

### 2.8 Caecum metabolomic analysis

Extraction of caecal water (n=10) was performed by mixing 50-100mg caecum content with water (MilliQ®, caecum content to water ratio 1:4) and centrifuging at 16000 x g for 10min at 4°C. Supernatant was transferred to a spinX centrifuge filter and centrifuged at 15000 x g for 5min at 4°C. Caecum metabolomics were conducted by MS-Omics (Denmark) using UHPLC-MS-MS. Sample analysis was carried out by MS-Omics as follows. The analysis was carried out using a Thermo Scientific Vanquish LC coupled to Thermo Q Exactive HF MS. An electrospray ionization interface was used as ionization source. Analysis was performed in negative and positive ionization mode. The UPLC was performed using a slightly modified version of the protocol described by Catalin *et al*. (UPLC/MS Monitoring of Water-Soluble Vitamin Bs in Cell Culture Media in Minutes, Water Application note 2011, 720004042en). Peak areas were extracted using Compound Discoverer 3.1 (Thermo Scientific). Identification of compounds were performed at four levels; Level 1: identification by retention times (compared against in-house authentic standards), accurate mass (with an accepted deviation of 3ppm), and MS/MS spectra, Level 2a: identification by retention times (compared against in-house authentic standards), accurate mass (with an accepted deviation of 3ppm). Level 2b: identification by accurate mass (with an accepted deviation of 3ppm), and MS/MS spectra, Level 3: identification by accurate mass alone (with an accepted deviation of 3ppm).

### 2.9 Bioinformatic analysis

Differential expression analyses were limited to non-drug-related features (metabolites) annotated at the highest confidence levels 1 and 2a (caecal features, n=212). All analyses were performed in R (version 4.1.1). Raw feature peak area values below their associated limit of detection (as reported by MsOmics) were considered missing (*i.e.*, set to “*NA*”), and only features with a maximum of 25% missingness per condition were retained for quantification (caecal features remaining, n=201). Additionally, to remove features displaying high technical variance, only metabolites with <10% relative standard deviation in the pooled quality control samples were retained (caecal features remaining, n=175). Data were subsequently normalized with variance stabilizing normalization (VSN), using the “*vsn*” package (allowing for 10% outliers, by default). While originally developed for microarray data (Huber *et al*., 2002), VSN has successfully been applied in untargeted (Li *et al*., 2016) and simulated (Jauhiainen *et al*., 2014) metabolomics of comparable dataset size to the one herein, capitalizing on similar mean-variance relationships (van den Berg *et al*., 2006) and error models (Rocke & Durbin, 2001). The generalised logarithm base 2 (glog_2_) transformation employed in VSN approximates the standard log_2_ function for values >>0. The “*limma*” package was used for differential expression analysis, with trend = TRUE and robust = TRUE in the eBayes function (Ritchie *et al*., 2015). Resulting feature *p*-values were adjusted for multiple comparisons with the Benjamini-Hochberg method, with a 5% false discovery rate (FDR) threshold for significance.

### 2.10 Statistical Analysis

Data were checked for outliers using the Grubbs outlier test, and for normality using Shapiro-Wilk. With the exception of running distance and Morris Water Maze training, data were analysed using two-way ANOVA in GraphPad Prism and SPSS. Running distance data were analysed using a two-way ANOVA with repeated measures, with post-hoc analysis comparing each week using Sidak’s multiple comparisons test. Morris Water Maze training outcomes were analysed using a two-way ANOVA with repeated measures, with post hoc analysis on individual training days using two-way ANOVA and Tukey’s multiple comparisons test. Post hoc analyses of all other two-way ANOVAs were performed using the Tukey’s multiple comparisons test. In the case of non-normally distributed data, data were analysed for differences in group distributions using Kruskal-Wallis with pairwise Mann-Whitney U post hoc comparisons. Statistical significance was set at p<0.05, with data presented as means ± standard error of the mean (SEM).

## 3. Results

### 3.1 Exercise attenuated cafeteria diet-induced increases in body weight gain and adipose tissue

There was a significant main effect of diet [F(1,92)=34.39, p<0.0001] and exercise [F(1,92)=16.61, p<0.0001] on body weight gain (Figure 1B), but no diet-exercise interaction. Post-hoc analysis revealed that CAF diet increased weight gain in sedentary [CAF-SED vs. CTRL-SED p<0.0001] as well as exercising [CAF-EX vs. CTRL-EX p<0.01] animals. Exercise showed a non-significant trend to reduce body weight gain in standard chow-fed animals [CTRL-EX vs. CTRL-SED p=0.071] and significantly reduced body weight gain in CAF diet-fed animals [CAF-EX vs. CAF-SED p<0.01].

Repeated measures ANOVA showed a significant effect of time [F(3.003,66.06)=50.67, p<0.0001], as well as a significant diet-time interaction [F(7,154)=2.43, p<0.05], on the average weekly running distance (Figure 1C). However, post-hoc analysis indicated no significant differences between the standard chow and CAF diet-fed groups in any given week.

There were significant main effects of diet [F(1,89)=84.04, p<0.0001] and exercise [F(1,89)=65.24, p<0.0001], as well as a significant diet-exercise interaction [F(1,89)=9.54, p<0.01] on the body weight-adjusted epidydimal white adipose tissue (eWAT) weight (Figure 1D). Post-hoc comparison revealed that a CAF diet increased eWAT weight in sedentary and exercising animals [CAF-SED vs. CTRL-SED p<0.0001; CAF-EX vs. CTRL-EX p<0.001]. On the other hand, exercise decreased eWAT weight in animals with access to standard chow [CTRL-EX vs. CTRL-SED p<0.01] and attenuated an increase in eWAT weight in animals with access to a CAF diet [CAF-EX vs. CAF-SED p<0.0001].

Furthermore, there were significant main effects of diet [F(1,91)=14.39, p<0.001] and exercise [F(1,91)=10.02, p<0.01], but not their combination, on the body weight-adjusted brown adipose tissue (BAT) weight (Figure 1E). Post-hoc testing showed that a CAF diet increased BAT weight in sedentary animals [CAF-SED vs. CTRL-SED p<0.01], but this effect was not present in exercising animals [CAF-EX vs. CAF-SED p<0.05].

### 3.2 Exercise exerted a modest anxiolytic effect irrespective of diet

Anxiety-like behaviour was assessed in the elevated plus maze (EPM) and novelty-suppressed feeding test (NSF). In the EPM, there was a trend for an exercise-induced increase in percent time spent in open arms [F(1,43)=3.69, p=0.062, Figure 2A]. There was no effect of a CAF diet on percent time spent in open arms, nor a diet-exercise interaction.

In the NSF, there was a significant difference between groups in the latency to eat [H(3)=10.74, p<0.05, Figure 2B]. Post-hoc analysis showed that exercise decreased the latency to eat in both standard chow-fed and CAF diet-fed groups, compared to sedentary controls, although this did not quite reach statistical significance [CTRL-EX vs. CTRL-SED p=0.056; CAF-EX vs. CAF-SED p=0.058]. After the NSF behavioural test, animals were given standard chow in the home cage for 30min, to control for differences in food interest which could in turn confound data obtained in the NSF test. There was a main effect of diet [F(1,42)=30.14, p<0.0001], but not of exercise, on body weight-adjusted standard chow consumption post testing (Figure 2C). In addition, there was a trend for a diet-exercise interaction, although this did not quite reach statistical significance [F(1,42)=3.70, p=0.061]. Post-hoc analysis revealed that a CAF diet significantly reduced post-test chow consumption in exercising animals [CAF-EX vs. CTRL-EX p<0.001], and non-significantly in sedentary animals [CAF-SED vs. CTRL-SED, p=0.071]. While this suggests that animals with access to a CAF diet were less interested in standard chow due to CAF food supplementation, it did not impact anxiety-like behaviour in the NSF. The lack of exercise effects on post-test chow consumption indicates that the reduced latency to eat was not due to increased interest in standard chow, but rather decreased anxiety-like behaviour.

### 3.3 Exercise mitigated a cafeteria diet-induced increase in immobility in the forced swim test

To measure antidepressant-like behaviour, animals underwent the forced swim test (FST) and their swimming, climbing and immobility scores were determined (Figure 2D). There was a significant main effect of diet [F(1,43)=10.41, p<0.01] and exercise [F(1,43)=7.87, p<0.01], but no diet-exercise interaction, on the immobility score (Figure 2D). Animals with access to a CAF diet showed increased immobility compared to standard chow-fed counterparts (CAF-SED vs. CTRL-SED p<0.05). This effect was mitigated by exercise (CAF-EX vs. CAF-SED p<0.05). There were significant main effects of a CAF diet [F(1,43)=8.80, p<0.001] and exercise [F(1,43)=4.39, p<0.05] on the swimming score (Figure 2D). However, post-hoc analysis did not reveal significant differences between groups, although there was a trend for a CAF diet to decrease the swimming score in exercising animals [CAF-EX vs. CTRL-EX p=0.060]. No effects of CAF diet or exercise were observed on climbing behaviour, although there was a non-significant trend for a diet-exercise interaction [F(1,43)=3.08, p=0.087, Figure 2D].

### 3.4 A cafeteria diet and exercise, alone or in combination, did not affect reward-seeking behaviour in the female urine sniffing test, or activity in the open field test

Animals were tested for reward-seeking behaviour in the female urine sniffing test (FUST). No effects of either diet or exercise, or a diet-exercise interaction, were observed for preference to sniff urine compared to water (Figure 2E). The OF test was used to assess general locomotion and anxiety-like behaviour. There were no significant effects of a CAF diet or exercise, or a diet-exercise interaction on total distance moved (Figure 2F) or time in centre (Figure 2G) in the test.

### 3.5 Neither a cafeteria diet nor exercise robustly altered pattern separation or recognition memory

Pattern separation was evaluated in the modified spontaneous location recognition (MSLR) test (Figure 3A). There were no effects of CAF diet or exercise on the discrimination ratio in the large separation (*i.e.*, pattern separation with low contextual overlap) test (Figure 3B). However, there were non-significant trends of diet [F(1,43)=3.69, p=0.061] and exercise [F(1,43)=3.98, p=0.052] on the discrimination ratio in the small separation (*i.e.*, pattern separation with high contextual overlap) test (Figure 3B), although post-hoc testing did not reveal significant differences between any of the groups.

**Figure 3:**
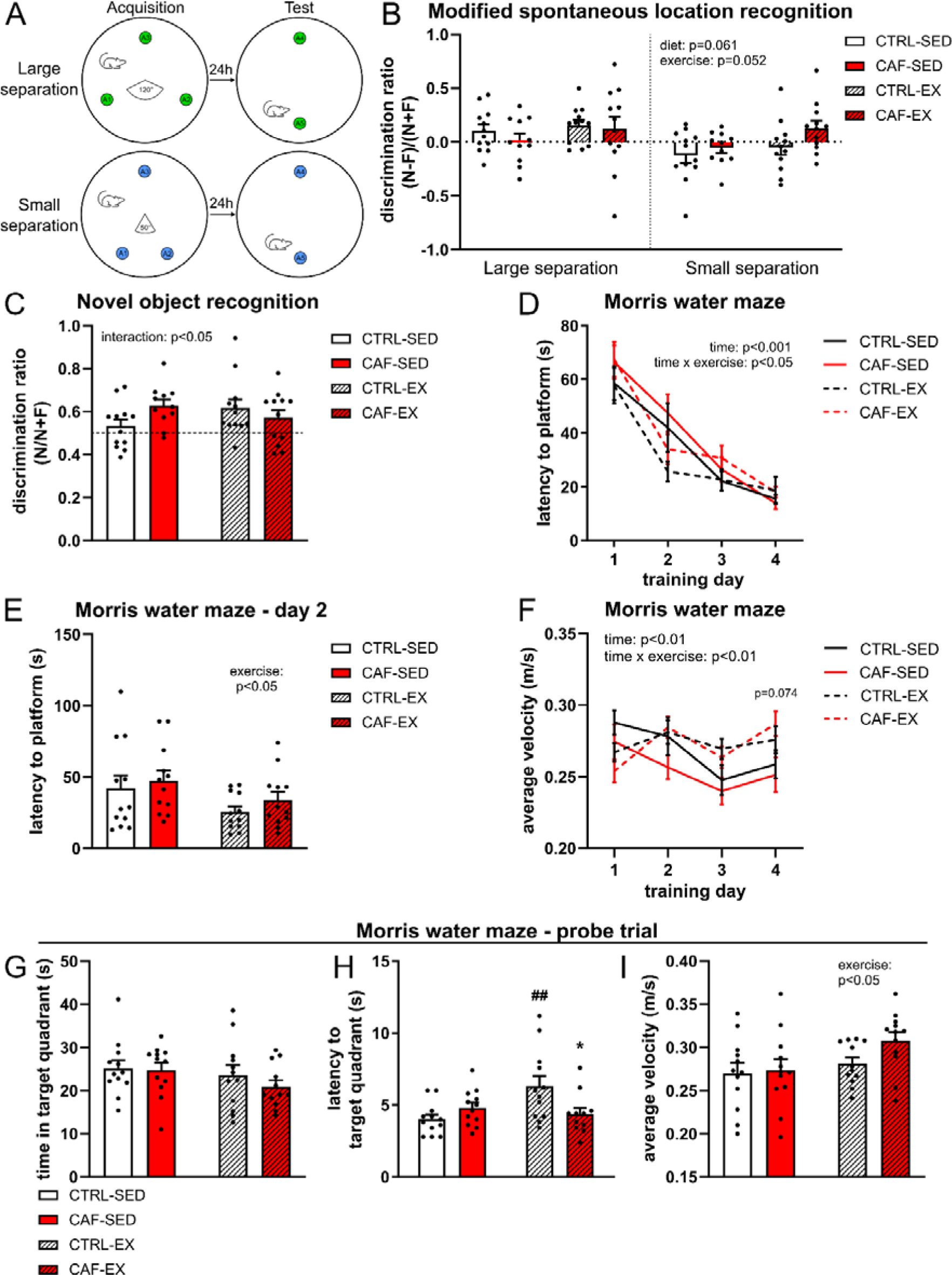
**(A)** Schematics of the large and small separation in the MSLR task. **(B)** Discrimination ratio in the large and small separation of the Modified Spontaneous Location Recognition (MSLR) ((novel (N) + familiar (F))/(N-F), n=12). **(C)** Discrimination ratio in the novel object recognition (NOR) task (N/(N+F), n=12). **(D, E)** Average latency (s) to find the platform in the Morris water maze (MWM), on **(D)** all training days and **(E)** training day 2 (n=12). **(F)** Average swimming velocity during MWM training (n=12). **(G)** Cumulative time spent in the target quadrant during the probe trial (n=12). **(H)** Average latency to reach the target quadrant during the probe trial (n=11-12). **(I)** Average swimming velocity during the probe trial (n=12). All data are expressed as mean ±SEM, *p<0.05 vs. corresponding treatment group with access to standard chow; ##p<0.01 vs. corresponding sedentary treatment group; p=0.074 CAF-EX vs. CAF-SED.

In the NOR test, which was used to assess recognition memory, there were no effects of CAF diet or exercise, although a significant diet-exercise interaction was observed [F(1,43)=4.17, p<0.05, Figure 3C]. However, post-hoc analysis revealed no significant differences between any of the groups.

### 3.6 Exercise showed a trend to improve spatial learning and search strategies, but not spatial memory

Spatial learning and memory were assessed in the Morris water maze (MWM). During training, there was a significant effect of time [F(3,24)=54.88, p<0.001], but not of a CAF diet or exercise individually or in combination, on the latency to find the platform (Figure 3D). However, there was a significant interaction of time and exercise [F(3,24)=3.80, p<0.05]. Two-way ANOVA analysis of the individual training days showed that on the second training day, there was a main effect of exercise on the latency to find the platform [F(1,44)=4.98, p<0.05, Figure 3E], but post-hoc analysis revealed no significant differences between the groups. There was a significant effect of time [F(3,30)=5.42, p<0.01], but not of a CAF diet or exercise, on the average swimming velocity during training, as well as a significant interaction of time and exercise [F(3,30)=5.77, p<0.01, Figure 3F]. Two-way ANOVA analysis of individual training days revealed a main effect of exercise on velocity the first [F(1,43)=5.13, p<0.05], third [F(1,44)=6.80, p<0.05], and fourth [F(1,44)=6.83, p<0.05] training day. Post-hoc analysis did not show significant differences in between groups, although there was a trend for exercise to increase swimming velocity in animals with access to a CAF diet on the fourth training day [CAF-EX vs. CAF-SED p=0.074].

During the probe trial, there were no effects of either diet, exercise, or their combination on the time spent in the target quadrant (Figure 3G), but there was a significant difference in group distributions of the latency to the first visit to target quadrant [H(3)=8.25, p<0.05, Figure 3H]. Post-hoc analysis indicated that exercise increased the latency in standard chow-fed, but not CAF diet-fed animals (CTRL-EX vs. CTRL-SED p<0.01; CAF-EX vs. CTRL-EX p<0.05). Two-way ANOVA analysis of the average velocity in the probe trial indicated a main effect of exercise [F(1,44)=4.51, p<0.05, Figure 3I], but not of a CAF diet or their combination. However, post-hoc analysis did not reveal significant differences between the groups.

Spatial search strategies in the MWM were also assessed (Figure 4). There was a significant effect of time [F(3,33)=74.00, p<0.001] but not of a CAF diet and/or exercise on the mean distance from platform (Figure 4A). However, exercise showed a significant interaction with time [F(3,33)=3.98, p<0.05]. There was a significant main effect of exercise [F(1,44)=7.04, p<0.05] on the mean distance from platform on the second training day (Figure 4B), but no significant differences between individual groups. There was a significant effect of time [F(3,33)=46.47, p<0.001] as well as a significant diet-exercise interaction [F(1,11)=5.13, p<0.05, Figure 4C] on the average cognitive score during training, which is a measure of search strategy whereby a higher score is indicative of more efficient and spatially precise searching. On the second training day (Figure 4D), there was a trend towards an effect of exercise [F(1,44)=3.30, p=0.076] and diet-exercise interaction [F(1,44)=3.00, p=0.090] although these did not reach statistical significance. There was no effect of CAF diet on the average cognitive score on the second training day (Figure 4D).

**Figure 4:**
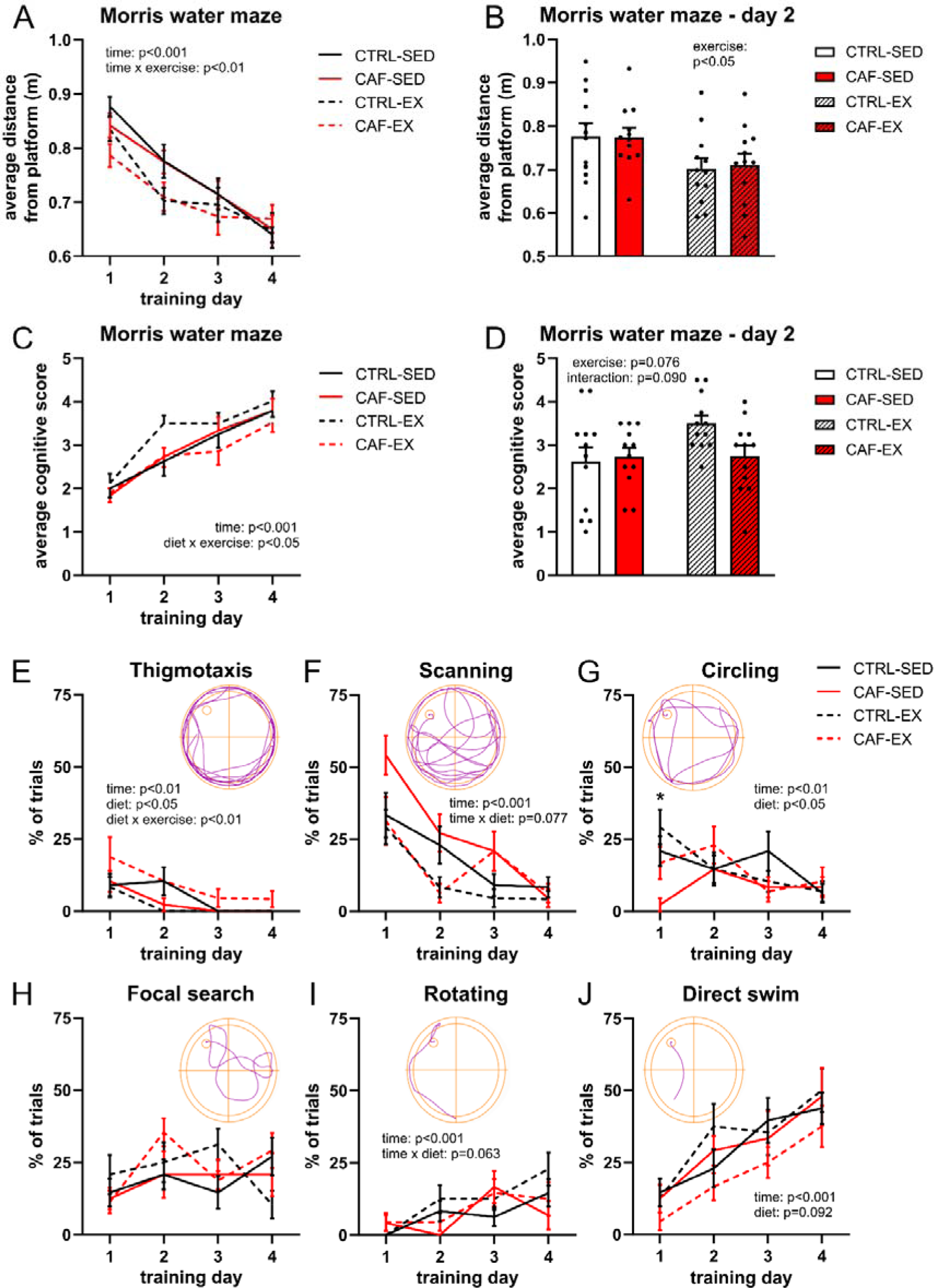
**(A, B)** Mean distance from the platform in the MWM on **(A)** all training days and **(B)** training day 2 (n=12). **(C, D)** Average cognitive score in the MWM on **(C)** all training days and **(D)** training day 2 (n=12). **(E-J)** Percentage of trials spent showing **(E)** thigmotaxis, **(F)** circling, **(G)** scanning, **(H)** focal search, **(I)** rotating, and **(J)** direct search behaviour on all training days (n=12). All data are expressed as mean ±SEM, *p<0.05 CAF-SED vs. CTRL-SED.

There were significant effects of time [F(3,21)=6.46, p<0.01] and CAF diet [F(1,7)=5.65, p<0.05], but not of exercise, on thigmotaxis behaviour, as well as a significant diet-exercise interaction [F(1,7)=14.00, p<0.01 Figure 4E]. However, Kruskal-Wallis analysis of individual training days did not reveal any significant differences between groups. In addition, there was a significant effect of time [F(3,24)=38.34, p<0.001], but not of a CAF diet or exercise, and a trend for a diet-time interaction [F(3,24)=2.58, p=0.077] on scanning behaviour (Figure 4F). There was a significant effect of time [F(3,24)=6.73, p<0.01] and diet [F(1,8)=9.31, p<0.05] but not of exercise or any interactions on circling behaviour (Figure 4G). Interestingly, there was a significant difference in group distributions of circling behaviour on the first training day [H(3)=11.85, p<0.01, Figure 4G]. Sedentary animals with access to a CAF diet showed decreased use of circling behaviour on training day one compared to animals with access to standard chow (CAF-SED vs. CTRL-SED p<0.05), which was not observed in exercising animals. There were no effects on focal searching (Figure 4H). There was a significant effect of time [F(3,21)=10.80, p<0.001], but not of a CAF diet or exercise, and a trend for a diet-time interaction [F(3,21)=2.84, p=0.063] on rotating behaviour (Figure 4I). There was a significant effect of time [F(3,30)=14.34, p<0.001], and a non-significant effect of a CAF diet [F(1,10)=3.49, p=0.092], on direct swims, but no effect of exercise (Figure 4J).

### 3.7 A cafeteria diet blunted an exercise-induced increase in adult hippocampal neurogenesis

The number of maturing hippocampal neurons was determined using immunohistochemical staining of doublecortin (DCX), a marker of immature neurons in the dentate gyrus. In the total dentate gyrus, there was a significant main effect of exercise [F(1,20)=17.48, p<0.001] but not diet on the number of DCX^+^ cells/mm^2^ (Figure 5A). In addition, there was a significant diet-exercise interaction [F(1,20)=7.39, p<0.05]. Post-hoc analysis revealed that in standard chow-fed animals, exercise significantly increased the number of DCX^+^ cells/mm^2^ [CTRL-EX vs. CTRL-SED p<0.001], but this effect was not observed in CAF diet-fed animals. Indeed, in exercising animals, a CAF diet showed a trend to decrease the number of DCX^+^ cells/mm^2^ [CAF-EX vs. CTRL-EX p=0.086].

**Figure 5:**
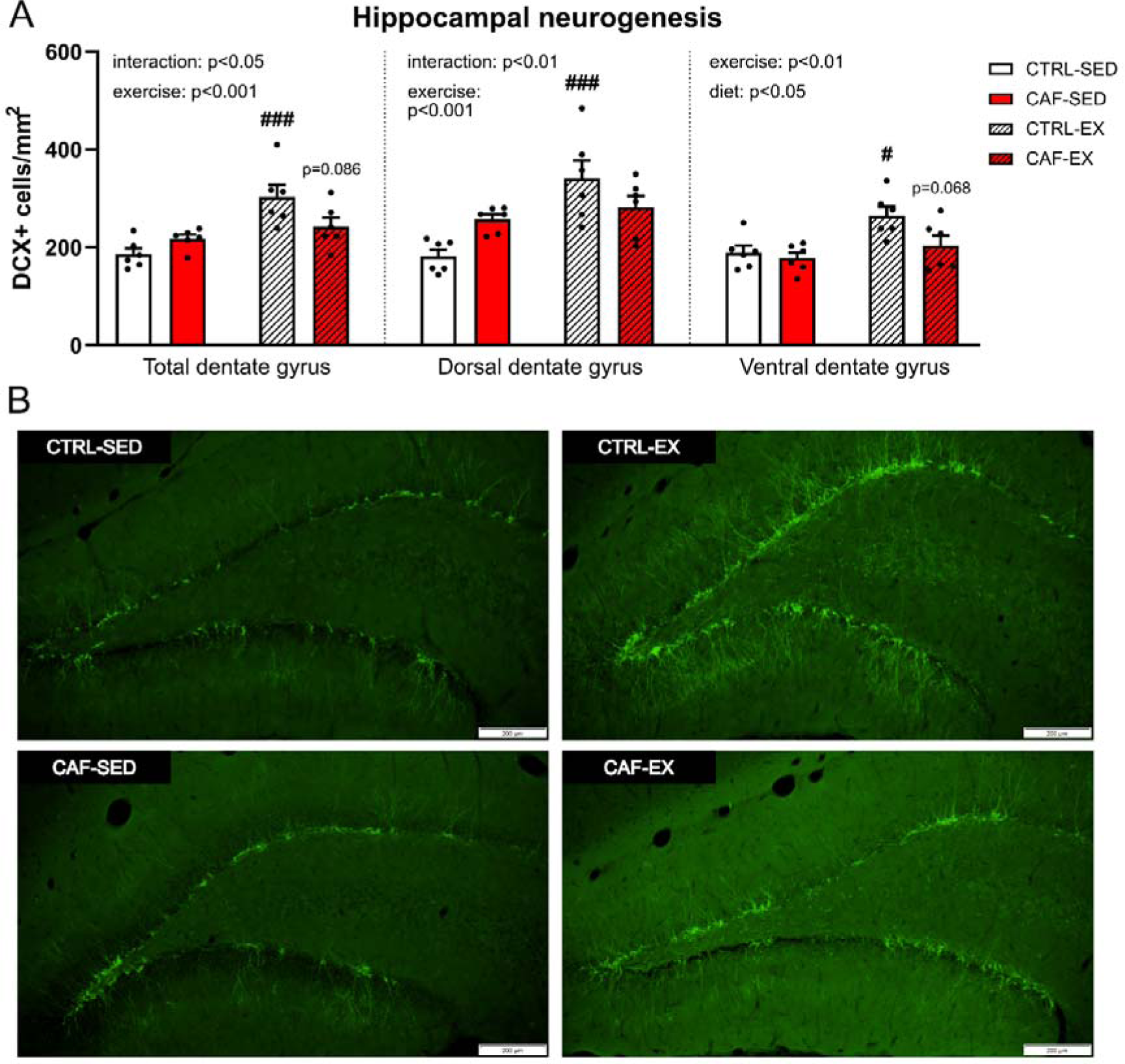
**(A)** Number of immature neurons in the dentate gyrus of the hippocampus (DCX^+^ cells/mm^2^, n=6). **(B)** Representative images of DCX neurons in dentate gyrus taken at 10x magnification. All data are expressed as mean ±SEM, p=0.086 & p=0.068 vs. corresponding treatment group with access to standard chow; #p<0.05 & ###p<0.001 vs. corresponding sedentary treatment group.

In the dorsal dentate gyrus, exercise similarly showed a significant main effect on the number of DCX^+^ cells/mm^2^ [F(1,20)=15.33, p<0.001, Figure 5A,B]. While there was no effect of diet, a significant interaction between diet and exercise was observed [F(1,20)=8.28, p<0.01]. Post-hoc analysis showed that exercise significantly increased the number of DCX^+^ cells/mm^2^ in standard chow-fed [CTRL-EX vs. CTRL-SED p<0.001] but not in CAF diet-fed animals.

In the ventral dentate gyrus, there was a significant main effect of exercise [F(1,20)=9.29, p<0.01] and CAF diet [F(1,20)=4.98, p<0.05], but no diet-exercise interaction on the number of DCX^+^ cells/mm^2^ (Figure 5A). Post-hoc analysis revealed that in standard chow-fed animals, exercise significantly increased the number of DCX^+^ cells/mm^2^ [CTRL-EX vs. CTRL-SED p<0.05], an effect which was not observed in CAF diet-fed animals. In addition, a CAF diet showed a trend to decrease the number of DCX^+^ cells/mm^2^ in exercising animals [CAF-EX vs. CTRL-EX p=0.068]. Representative images of DCX-positive cells in dorsal hippocampus are shown in Figure 5B.

### 3.8 A cafeteria diet blunted an exercise-induced increase in total GLP-1 plasma concentrations, while exercise increased PYY and attenuated cafeteria diet-induced increases in insulin and leptin

Changes in metabolic hormones such as glucagon-like peptide (GLP) 1, insulin, and leptin have previously been linked with depression, anxiety, cognitive impairment, and altered neurogenesis (Isacson *et al*., 2011; Zou *et al*., 2019; Spinelli *et al*., 2019). Therefore, we measured the effects of CAF diet and exercise on plasma concentrations of metabolic hormones in animals from both experiments. Kruskal-Wallis testing of total GLP-1 (tGLP-1) concentrations revealed significant differences between groups [H(3)=12.61, p<0.01, Figure 6A], with post-hoc analysis showing that exercise increased tGLP-1 concentrations in standard chow-fed animals [CTRL-EX vs. CTRL-SED p<0.01]. A CAF diet did not alter tGLP-1 concentrations in sedentary animals [CAF-SED vs. CTRL-SED p>0.05] but prevented the exercise-induced increase in tGLP-1 [CAF-EX vs. CTRL-EX p<0.05].

**Figure 6:**
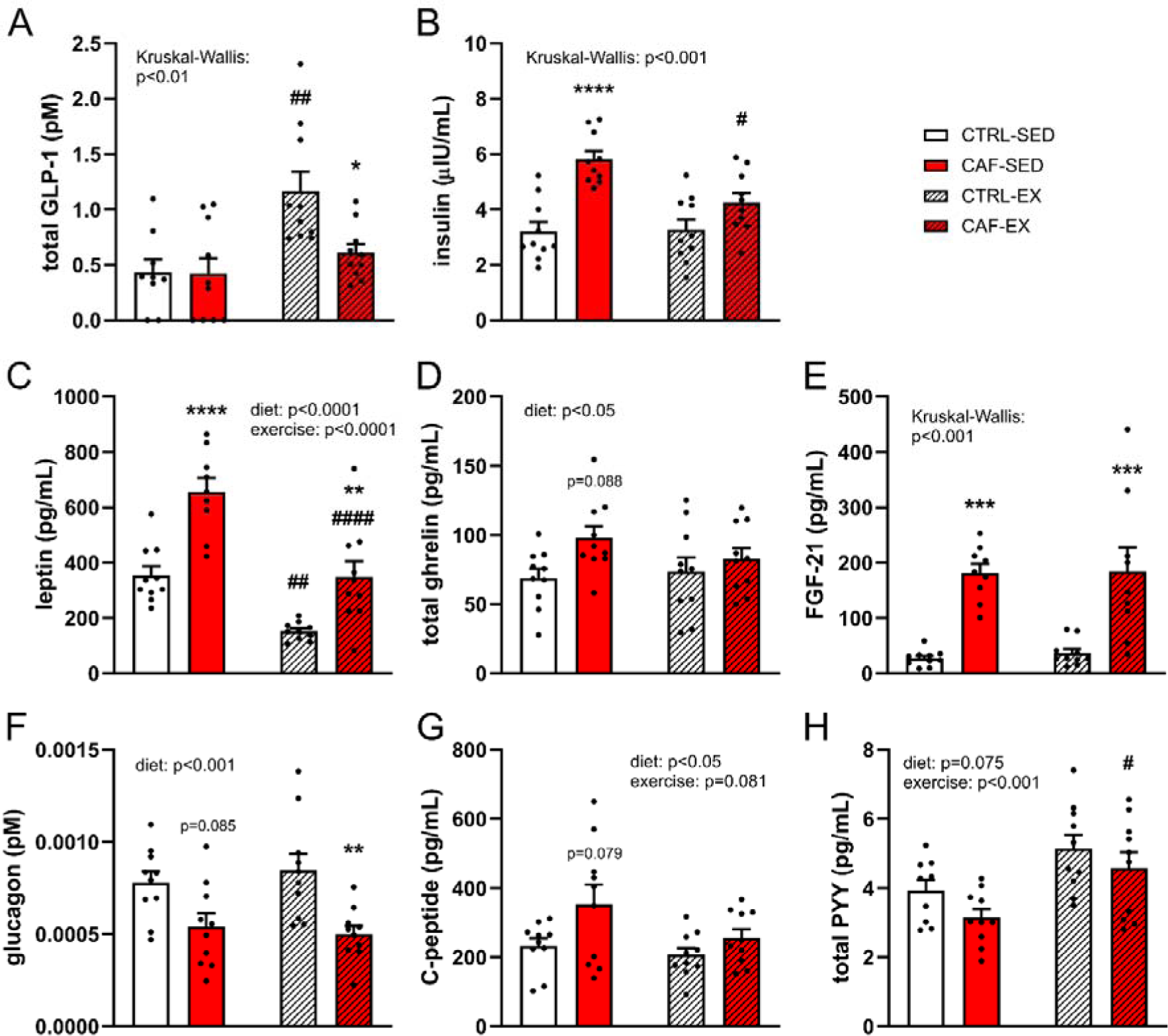
Concentrations of plasma metabolic hormones (n=9-10) **(A)** total GLP-1 (pM), **(B)** insulin (μIU/mL), **(C)** leptin (pg/mL), **(D)** total ghrelin (pg/mL), **(E)** FGF-21 (pg/mL), **(F)** glucagon (pM), **(G)** C-peptide (pg/mL) and **(H)** total PYY (pg/mL). All data are expressed as mean ±SEM, p=0.088, p=0.085, p=0.079, *p<0.05, **p<0.01, ***p<0.001 & ****p<0.0001 vs. corresponding treatment group with access to standard chow; #p<0.05, ##p<0.01 & ####p<0.0001 vs. corresponding sedentary treatment group.

Plasma insulin concentrations showed significant differences in group distributions [H(3)=18.99, p<0.001, Figure 6B]. Post-hoc analysis revealed that in sedentary animals, a CAF diet increased insulin concentrations [CAF-SED vs. CTRL-SED p<0.001]. Exercise did not alter insulin concentrations in animals with access to standard chow [CTRL-EX vs. CTRL-SED p>0.05] but attenuated the CAF diet-induced increase in insulin [CAF-EX vs. CAF-SED p<0.05].

There were significant main effects of both CAF diet [F(1,35)=36.11, p<0.0001, Figure 6C] and exercise [F(1,35)=38.01, p<0.0001], but no diet-exercise interaction on plasma leptin concentrations. Post-hoc analysis showed that a CAF diet significantly increased leptin in both sedentary and exercising animals [CAF-SED vs. CTRL-SED p<0.0001; CAF-EX vs. CTRL-EX p<0.01], and that exercise decreased leptin concentrations irrespective of diet [CTRL-EX vs. CTRL-SED p<0.01; CAF-EX vs. CAF-SED p<0.0001], leading to a normalisation of leptin concentrations in exercising animals with access to a CAF diet [CAF-EX vs. CTRL-SED p>0.05].

A significant main effect of diet [F(1,36)=5.11, p<0.05, Figure 6D], but not exercise was observed on plasma total ghrelin concentrations. No diet-exercise interaction was found. Post-hoc analysis did not indicate any significant differences between groups, although there was a non-significant trend for a CAF diet to increase total ghrelin in sedentary animals [CAF-SED vs. CTRL-SED p=0.088].

Plasma FGF-21 concentration distributions were significantly different across groups [H(3)=25.51, p<0.001, Figure 6E], with post-hoc testing indicating that a CAF diet increased FGF-21 concentrations in both sedentary and exercising animals [CAF-SED vs. CTRL-SED p<0.001; CAF-EX vs. CTRL-EX p<0.01].

There was a significant main effect of diet on plasma glucagon concentrations [F(1,36)=18.09, p<0.001, Figure 6F], but no effect of exercise or a diet-exercise interaction. Post-hoc analysis showed that a CAF diet significantly decreased glucagon concentrations in exercising animals [CAF-EX vs. CTRL-EX p<0.01] and non-significantly in sedentary animals [CAF-SED vs. CTRL-SED p=0.085].

There was a significant main effect of diet [F(1,36)=6.15, p<0.05] on the plasma concentration of C-peptide (Figure 6G). There was a trend for an effect of exercise but this did not reach statistical significance [F(1,36)=3.22, p=0.081, Figure 6G]. There was no interaction between diet and exercise. No significant differences between groups were revealed after post-hoc analysis, although there was a trend for a CAF diet to increase C-peptide concentrations in sedentary animals although this did not reach statistical significance [CAF-SED vs. CTRL-SED p=0.079].

Lastly, there was a trend for an effect of CAF diet [F(1,35)=3.36, p=0.075, Figure 6H] and a significant main effect of exercise [F(1,35)=13.24, p<0.001], but no diet-exercise interaction on plasma total PYY concentrations. Post-hoc analysis indicated that exercise significantly increased PYY in CAF diet-fed, but not standard chow-fed animals [CAF-EX vs CAF-SED p<0.05].

### 3.9 Exercise attenuated a cafeteria diet-induced decrease in caecal metabolites anserine, indole-3-carboxylate, and deoxyinosine

There were significant main effects of both a CAF diet [F(1,44)=73.80, p<0.0001] and exercise [F(1,44)=43.98, p<0.0001] on body weight-adjusted caecum weight, but no interaction was observed (Figure 7A). Post-hoc testing revealed that a CAF diet significantly reduced caecum weight in sedentary and exercising animals [CAF-SED vs. CTRL-SED p<0.0001; CAF-EX vs. CTRL-EX p<0.0001]. Conversely, exercise increased caecum weight irrespective of diet [CTRL-EX vs. CTRL-SED p<0.001; CAF-EX vs. CAF-SED p<0.001].

**Figure 7:**
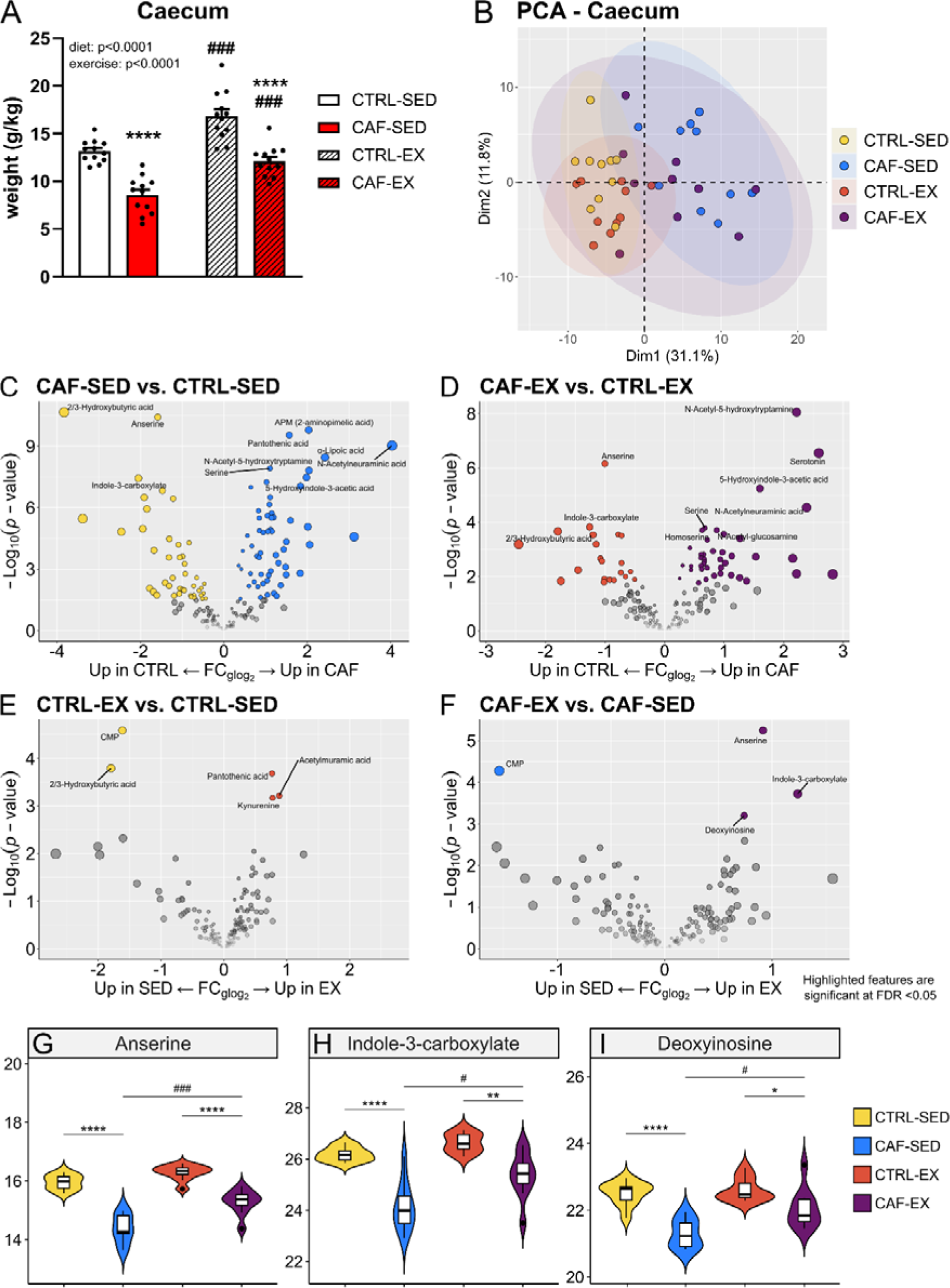
**(A)** Effects of a CAF diet and exercise on body weight-adjusted caecum weight (g/kg, n=12). Data are expressed as mean ±SEM, ****p<0.0001 vs. corresponding treatment group with access to standard chow; ###p<0.001 vs. corresponding sedentary treatment group. **(B)** Principal component analysis (PCA) with 95% concentration ellipses showing effects of a CAF diet and exercise on caecal metabolomes (n=10) **(C-F)** Volcano plots of quantified caecal metabolites (n=175 features), comparing **(C)** a cafeteria diet (CAF) vs. standard chow (CTRL) in sedentary (SED) animals, **(D)** CAF vs. CTRL in exercising (EX) animals, **(E)** EX vs. SED in CTRL diet-fed animals, and **(F)** EX vs. SED in CAF diet-fed animals. Highlighted features represent upregulated metabolites (p-value adjusted for false discovery rate (FDR) <0.05). **(G-I)** Violin and box- and whisker plot (median represented by horizontal line) of the normalised peak area of caecal metabolites **(G)** anserine, **(H)** indole-3-carboxylate, and **(I)** deoxyinosine (all n=10, FDR-adjusted p-values; *p<0.05, **p<0.01, & ****p<0.0001 vs. corresponding standard chow-fed treatment group; p=0.094, #p<0.05, & ###p<0.001 vs. corresponding sedentary treatment group). Full list of quantified metabolites available in Supplemental Table 2. Abbreviations: fold change (FC), generalised logarithm base 2 (glog_2_).

To identify gut-derived metabolites that might be potentially associated with alterations in behaviour and AHN, untargeted metabolomic analysis of caecal content was performed. Principal component analysis suggested an effect particularly of a CAF diet on the caecal metabolome (Figure 7B). Differential expression analyses revealed that in caecal content of sedentary animals, a CAF diet induced differential expression (FDR-adjusted p<0.05) of 100 out of 175 metabolites (Figure 7C), while in exercising animals, a CAF diet induced differential expression of 62 out of 175 metabolites (Figure 7D), compared to standard chow. On the other hand, voluntary running exercise induced differential expression of 5 out of 175 metabolites in standard chow-fed animals (Figure 7E), and 4 out of 175 metabolites in CAF diet-fed animals (Figure 7F), compared to sedentary controls. Supplemental Table 2 provides a complete list of quantified features.

The top 10 differentially expressed (FDR-adjusted p<0.05) features within each of the group contrasts were further examined (Supplementary Table 2), and metabolites that had a significant diet-exercise interaction are shown in Figure 7G-I. The abundance of anserine was decreased by a CAF diet in sedentary as well as exercising animals [CAF-SED vs. CTRL-SED p<0.0001; CAF-EX vs. CTRL-EX p<0.0001]. Exercise attenuated this CAF diet-induced decrease in anserine abundance [CAF-EX vs. CAF-SED p<0.001, Figure 7G].

Similar to anserine, a CAF diet decreased the abundance of indole-3-carboxylate in sedentary and exercising animals [CAF-SED vs. CTRL-SED p<0.0001; CAF-EX vs. CTRL-EX p<0.01], and exercise attenuated a CAF diet-induced decrease in indole-3-carboxylate abundance [CAF-EX vs. CAF-SED p<0.05, Figure 7H].

Likewise, the abundance of deoxyinosine was decreased by a CAF diet in both sedentary and exercising animals [CAF-SED vs. CTRL-SED p<0.0001; CAF-EX vs. CTRL-EX p<0.05], and exercise attenuated this CAF diet-induced decrease in deoxyinosine abundance [CAF-EX vs. CAF-SED p<0.05, Figure 7I].

A CAF diet increased the abundance of n-acetylneuraminic acid in sedentary as well as in exercising animals [CAF-SED vs. CTRL-SED p<0.0001; CAF-EX vs. CTRL-EX p<0.01]. There were no significant effects of exercise, although there was a trend for exercise to attenuate the CAF diet-induced increase in n-acetylneuraminic acid abundance [CAF-EX vs. CAF-SED p<0.094, Supplementary Table 2].

The abundance of 2/3-hydroxybutyric acid was decreased by a CAF diet in both sedentary and exercising animals [CAF-SED vs. CTRL-SED p<0.0001; CAF-EX vs. CTRL-EX p<0.01]. In addition, exercise decreased the abundance of 2/3-hydroxybutyric acid in animals with access to standard chow [CTRL-EX vs. CTRL-SED p<0.05], but not a CAF diet (Supplementary Table 2).

On the other hand, a CAF diet increased the abundance of pantothenic acid in sedentary and exercising animals [CAF-SED vs. CTRL-SED p<0.0001; CAF-EX vs. CTRL-EX p<0.01]. Exercise increased the abundance of pantothenic acid in animals with access to standard chow [CTRL-EX vs. CTRL-SED p<0.05], but not a CAF diet (Supplementary Table 2).

The abundance of kynurenine was increased by a CAF diet in sedentary [CAF-SED vs. CTRL-SED p<0.05] but not exercising animals, and by exercise in animals with access to standard chow [CTRL-EX vs. CTRL-SED p<0.05] but not a CAF diet (Supplementary Table 2). Similar effects were observed for acetylmuramic acid [CAF-SED vs. CTRL-SED p<0.0001; CTRL-EX vs. CTRL-SED p<0.05, Supplementary Table 2].

A CAF diet increased the abundance of 2-aminopimelic acid, α-lipoic acid, homoserine, n-acetyl-glucosamine, serotonin, serine, 5-hydroxyindole-3-acetic acid, and n-acetyl-5-hydroxytryptamine in both sedentary [CAF-SED vs. CTRL-SED all p<0.0001, Supplementary Table 2] and exercising animals [CAF-EX vs. CTRL-EX: 2-aminopimelic acid, p<0.01; α-lipoic acid, p<0.05; homoserine, p<0.01; n-acetyl-glucosamine p<0.01; serotonin, p<0.0001; serine, p<0.01; 5-hydroxyindole-3-acetic acid, p<0.001; n-acetyl-5-hydroxytryptamine, p<0.0001]. There was no effect of exercise on these metabolites.

Lastly, exercise decreased the abundance of CMP in animals with access to standard chow [CTRL-EX vs. CTRL-SED p<0.01] and a CAF diet [CAF-EX vs. CAF-SED p<0.01], but there was no effect of a CAF diet on this metabolite (Supplementary Table 2).

## 4. Discussion

This study investigated whether voluntary running exercise in young adult male Sprague-Dawley rats could mitigate negative effects of a CAF diet on anxiety-like, depression-like, and cognitive behaviours, and on AHN and metabolic hormones associated with these effects. Since the MGBA influences AHN and related behaviours in response to diet and exercise (Guzzetta *et al*., 2022; Nicolas *et al*., 2024), whether exercise could also alter effects of a CAF diet on gut microbial metabolites was also investigated. We demonstrated that a CAF diet increased immobility in the forced swim test (FST) and this was attenuated by exercise, suggesting that exercise exerted antidepressant-like effects in CAF diet-fed animals. While a CAF diet did not alter anxiety-like behaviour, exercise had moderate anxiolytic effects in the elevated plus maze (EPM) and novelty suppressed feeding (NSF) tests. No effects of either a CAF diet or exercise were observed on hedonic behaviour in the female urine sniffing test (FUST). Exercise exerted mild improvements in spatial learning in the MWM. A CAF diet blunted exercise-induced increases in AHN and plasma total GLP-1. On the other hand, exercise increased PYY and attenuated CAF diet-induced increases in insulin and leptin concentrations but did not mitigate a CAF diet-induced increase in FGF-21 and decrease in glucagon. A CAF diet tended to increase C-peptide and total ghrelin in sedentary but not exercising animals. Finally, exercise attenuated a CAF diet-induced reduction of the metabolites anserine, indole-3-carboxylate, and deoxyinosine in the caecum, an interesting finding given previous reports that these metabolites may play a role in memory or depression-like behaviour (Su *et al*., 2011; Caruso *et al*., 2021; Lu *et al*., 2022).

The effects of a CAF diet on adult rodent anxiety-like and depression-like behaviours have not been extensively investigated. One recent study reported that a two-week CAF diet intervention increased anxiety-like behaviour in adult male Wistar rats (Maciel Reis *et al*., 2023). In contrast, CAF diet did not induce anxiety-like behaviour in the present study. This might be explained by the experimental differences in strain, duration of CAF diet, or discrepancies in the CAF diet feeding protocols. Similar to our findings, a study in adolescent (6-week-old) and middle-aged (12-month-old) male Sprague-Dawley rats did not demonstrate robust effects of a 6-week CAF diet on anxiety-like behaviour (Warneke *et al*., 2014). In the present study, a CAF diet did not induce anhedonia as measured in the FUST. Likewise, a recent study reported that in male and female Wistar rats, a 3-week CAF diet initiated at weaning did not affect reward-seeking behaviour in the sucrose preference test, even though male animals with access to a CAF diet ingested less sucrose solution (Mota-Ramírez & Escobar, 2023). On the other hand, it was recently shown that in male Wistar rats, access to a CAF diet for 17 weeks from birth decreased sucrose preference, indicating anhedonia (Gonçalves *et al*., 2021). While that finding could indicate that the length of CAF diet feeding in the current study was not sufficient to induce anhedonic-like behaviour, it should also be acknowledged that exposure to sweet foods could alter the hedonic value of sucrose, thereby interfering with the sucrose preference test. Finally, several studies in adult male C57BL/6 mice have reported that varying lengths (3 weeks - 9 months) of a HFD during adulthood increased immobility in the FST (Tsai *et al*., 2018; Vagena *et al*., 2019; Seguella *et al*., 2021; Zhuang *et al*., 2022), supporting our findings that a CAF diet increases immobility in the FST in adult male rats.

Currently, little is known about the impact of a CAF diet on pattern separation, novel object recognition, and spatial learning and memory. Although it was reported that a 2-month or 20-week CAF diet intervention in adult male rats impaired recognition memory (Lewis *et al*., 2019; Bondan *et al*., 2019), this was not the case in the present study. Furthermore, when a CAF diet was administered during adolescence and later discontinued, impairments in novel object recognition memory were not observed during adulthood, despite a persistent increase in insulin and leptin concentrations (Nicolas *et al*., 2022), suggesting age-related interaction between metabolic hormones and cognitive behaviour. Lewis and colleagues demonstrated that in adult male Sprague-Dawley rats, a 20-week CAF diet decreased time spent in the target quadrant during the probe trial of the MWM, suggesting spatial memory impairment (Lewis *et al*., 2019). We did not observe this impairment although there was a CAF diet-induced decrease in circling behaviour on the first training day, which is indicative of altered search strategy. In fact, although it did not reach statistical significance, there were modest improvements rather than impairments in pattern separation with high contextual overlap in rats with access to a CAF diet. Importantly, the CAF food items used by Bondan *et al*. (2019) and Lewis *et al*. (2019) did not vary day-to-day. This lack of variety in dietary items may have contributed to the cognitive impairment observed, as it has been shown by others that environmental enrichment, which a daily variation in CAF diet may provide, improves performance in the NOR test (Mesa-Gresa *et al*., 2013). It is therefore possible that environmental enrichment through a varying diet could improve pattern separation as well, and conversely, a lack of variation might lead to stronger negative effects of a CAF diet on cognitive function.

The effects of exercise on anxiety-like, depression-like and cognitive behaviours have been more widely investigated than the impact of a CAF diet. In the present study, voluntary wheel running significantly decreased the latency to eat in the NSF test and tended to increase time spent in the open arms of the EPM, suggesting an anxiolytic effect of exercise which is in agreement with previous studies conducted in adult male mice in the EPM (Duman *et al*., 2008; Morgan *et al*., 2018). In contrast however, Fuss and colleagues reported an exercise-induced increase in anxiety-like behaviour in the EPM in adult male C57BL/6 mice, although these results may have been confounded by an exercise-induced reduction in general locomotor activity (Fuss *et al*., 2010). We did not observe an antidepressant-like effect of exercise alone in the FST or in the FUST in animals with access to standard chow. This finding in the FST agrees with other studies using longer exercise interventions (1.5-11 months) in adult male mice (Fuss *et al*., 2010; Morgan *et al*., 2018), but not with studies using 3-4-week interventions which demonstrated a decrease in immobility following exercise (Duman *et al*., 2008; Cunha *et al*., 2013). Our finding in the FUST that there were no effects of exercise on reward-seeking behaviour has similarly been reported by others in adult male rodents using the sucrose preference test (Sigwalt *et al*., 2011; Gilak-Dalasm *et al*., 2021).

While many studies have investigated the therapeutic effects of exercise during aging or in rodent models of Alzheimer’s disease (Caruso *et al*., 2024), in which cognitive function is impaired, a more limited number of studies have investigated effects of exercise on cognitive behaviours in healthy young adult male animals. We found that while there were no beneficial effects of wheel running on performance in the probe trial of the MWM, exercising animals with access to standard chow displayed modest improvements in spatial learning and search strategy on the second training day. Likewise, four weeks of either voluntary or treadmill running exercise has been reported to improve spatial learning and memory in adult male BALB/c mice (Liu *et al*., 2009). In the present study, exercise did not enhance pattern separation or recognition memory. This contrasts with a previous study reporting that 10-11 weeks of voluntary running exercise improved pattern separation in adult male C57BL/6 mice (Creer *et al*., 2010). It is possible that methodological differences in species/strain as well as duration/type of exercise intervention may explain this discrepancy. Given the dearth of studies, further research will be required to make firm conclusions about the impacts of exercise on cognitive function in young adult rats.

A key aim of the current study was to determine if exercise could attenuate any negative effects of a CAF diet on depression- and anxiety-like behaviours, as well as cognitive function. We found that increased immobility in the FST induced by a CAF diet was attenuated by exercise, suggesting an antidepressant-like effect of exercise in CAF diet animals. In the MWM, we found that a CAF diet-induced decrease in circling behaviour on the first training day of the MWM, indicating an alteration in search strategy, was not observed in exercising animals, suggesting a modest beneficial effect of exercise on search strategy in CAF diet-fed animals. In support, a beneficial effect of 23 weeks of treadmill exercise prevented HFD-induced impairments in MWM spatial learning in young adult male mice (Han *et al*., 2019), although notably the diet and exercise interventions were initiated during late adolescence. On the other hand, Sack and colleagues (2017) reported that in the puzzle box test, which is hippocampus-dependent, young adult male mice with access to a CAF diet displayed mild long term memory impairment, which was not attenuated by exercise. This suggests that the impact of exercise on cognitive impairments induced by Western-style diets may be context dependent.

Given previous reports of dietary and exercise modulation of AHN (Hueston *et al*., 2017), and the role of AHN in cognitive function and antidepressant action (Zhang *et al*., 2008; Clelland *et al*., 2009; O’Leary & Cryan, 2014), we investigated the effects of a CAF diet and voluntary wheel running on AHN. One study in juvenile Wistar rats indicated that a 12-week CAF diet provided from adolescence decreased the number of hippocampal DCX^+^ neurons in adulthood (Ferreira *et al*., 2018). However, these effects were not observed in the current study in adult animals, suggesting that a CAF diet given during adolescence rather than in adulthood may have a greater impact on AHN. On the other hand, we found that adult-initiated exercise increased the number of hippocampal DCX^+^ neurons in animals with access to standard chow, a finding that is in agreement with other studies in adult male rats and mice (Fuss *et al*., 2010; Nokia *et al*., 2016).

Interestingly, we found that a CAF diet blunted the effects of voluntary running exercise on the production of immature neurons, *i.e.*, DCX^+^ cells. In agreement, it was reported that in male C57BL/6 mice, treadmill exercise initiated during adolescence mitigated concurrent HFD-induced decreases in hippocampal cell proliferation and neuronal maturation in adulthood (Han *et al*., 2019). Contrary to the present findings however, it was reported that a CAF diet intervention had no effect on 9-10 weeks of voluntary wheel running exercise-induced increases in hippocampal DCX^+^ cells in adult male C57BL/6 mice (Sack *et al*., 2017). However, it is important to note that in contrast to the current study in adult male rats, Sack *et al*. found that the mice with access to a CAF diet had an increased average daily running distance compared to standard chow-fed animals. The increased amount of exercise could potentially have protected against negative effects of the CAF diet on AHN in these animals. In addition, the CAF diet used by Sack *et al*. (2017) varied from the one used in the present study in feeding frequency, with new food items given every 2-3 days rather than daily, which may also account for some discrepancies along with species differences.

Previous research has shown that increasing AHN improves pattern separation (Sahay *et al*., 2011). However, despite an exercise-induced increase in AHN, which was blunted by a CAF diet, there were no alterations on pattern separation ability in the MSLR test. Importantly, a recent study showed that the small separation task, which has a high contextual overlap, in the modified spontaneous location recognition test may not be sensitive enough to show effects of interventions that increase AHN. Instead, it was suggested that improvements may only be observed with a very high contextual overlap, using an extra small separation (Reichelt *et al*., 2021), thus offering a potential explanation for the lack of exercise-induced improvements in pattern separation in the MSLR despite the exercise-induced increase in AHN that was observed in the current study.

As depression, anxiety and cognitive impairment have been associated with metabolic alterations (Hryhorczuk *et al*., 2013; Perini *et al*., 2019), we measured circulating concentrations of metabolic hormones in animals from both experiments. In contrast to another study (Virtuoso *et al*., 2018), a CAF diet did not decrease tGLP-1 in sedentary animals in the current study, potentially due to differences in length of dietary intervention. We further report that exercise alone increased plasma tGLP-1 concentrations, which agrees with human studies where a single bout as well as 3 months of exercise increased circulating tGLP-1 (Martins *et al*., 2007; Hazell *et al*., 2017; Ataeinosrat *et al*., 2022). Interestingly, in the present study a CAF diet prevented an exercise-induced increase in total GLP-1. Previous research has shown that in male Sprague-Dawley rats (age unspecified), activation of the GLP-1 receptor using the agonist exendin-4 increased AHN and improved spatial memory in the radial arm maze (Isacson *et al*., 2011). Conversely, treatment with a GLP-1 receptor antagonist in adult male Wistar rats has been reported to attenuate an exercise-induced improvement in spatial learning in the MWM (Taati *et al*., 2022). Therefore, it is possible that CAF diet attenuation of the exercise-induced increase in AHN observed here is mediated by circulating concentrations of total GLP-1, although this was not accompanied by similar alterations in spatial learning or memory.

It is well-established that a CAF diet increases the concentrations of the metabolic hormones insulin and leptin (De Schepper *et al*., 1998; MacEdo *et al*., 2012; Higa *et al*., 2014*a*). In agreement, we found that a CAF diet increased plasma insulin and leptin concentrations in sedentary animals, but this was attenuated by exercise. This is further supported by previous research which showed that 8 weeks of treadmill exercise prevented CAF diet-induced increases in leptin and insulin in adult male C56BL/6J mice (Higa *et al*., 2014*b*). While leptin and insulin are known to increase AHN and related cognitive behaviours (Zou *et al*., 2019; Spinelli *et al*., 2019; Calió *et al*., 2021), resistance to these hormones due to prolonged elevation has been associated with depression-like behaviour and cognitive impairment (Van Doorn *et al*., 2017; Spinelli *et al*., 2019). Leptin resistance (induced by treatment with a leptin receptor antagonist) and hippocampal insulin resistance have previously been shown to increase immobility in the FST in adult male Sprague-Dawley rats (Macht *et al*., 2017; Reagan *et al*., 2021). Therefore, attenuation of CAF diet-induced increases in insulin and leptin by exercise may have contributed to the mitigating effects of exercise on CAF diet-induced immobility in the FST in the present study. Furthermore, given that white adipose tissue is known to release leptin and regulate insulin sensitivity (Guerre-Millo, 2002; Smith & Kahn, 2016), the exercise-induced decrease in adipose tissue in animals with access to a CAF diet observed in our study possibly underlies mitigating effects of exercise on a CAF diet-induced increase in leptin, insulin, and immobility in the FST.

In agreement with human studies, exercise increased plasma metabolic hormone PYY, (Martins *et al*., 2007; Broom *et al*., 2009; Cooper *et al*., 2011) and particularly in CAF diet-fed animals. Increased anxiety-like behaviour in the EPM and immobility in the FST was previously reported in PYY knockout mice (Painsipp *et al*., 2009), and centrally administered PYY has been shown to decrease anxiety-like behaviour in the EPM in adult male rats (Broqua *et al*., 1995). Therefore, the exercise-induced increase in PYY potentially contributed to its anxiolytic effects observed here.

Caecal metabolite differential expression was significantly altered by a CAF diet, which was attenuated by exercise. Specifically, exercise attenuated a CAF diet-induced decrease in caecal abundance of the metabolite anserine, supplementation with which has previously been associated with improved cognitive function and protection against Alzheimer’s disease (Caruso *et al*., 2021). For example, 8-week dietary supplementation of anserine in a mouse model of Alzheimer’s disease attenuated spatial memory deficits in the MWM and density of activated inflammatory astrocytes in the dentate gyrus (Kaneko *et al*., 2017). In elderly individuals without cognitive impairment, a 13-week supplementation with anserine and a related metabolite, carnosine, significantly improved cognitive function in the Mini Mental State Examination and the Short Test of Mental Status tests compared to placebo (Szcześniak *et al*., 2014). Given the role of anserine in modulating hippocampal cognitive function, future studies are needed to examine its impact on AHN and to determine if dietary supplementation with anserine could potentially counteract the negative effects of a CAF diet on exercise-induced increases in AHN.

Exercise also attenuated a CAF diet-induced decrease in the caecal abundance of deoxyinosine and indole-3-carboxylate. Interestingly, it has been reported that the hippocampal concentration of deoxyinosine, a nucleoside, was decreased in a mouse model of depression (Lu *et al*., 2022), whereas chronic variable stress has been shown to increase urine concentrations of indole-3-carboxylate (Su *et al*., 2011). Importantly, indole-3-carboxylate is biochemically related to the essential amino acid tryptophan, a precursor of the neurotransmitter serotonin which has been reported to negatively correlate with depression scores (Delgado, 1990; Gertsman *et al*., 2015). It is possible that the attenuation of a CAF diet-induced decrease in deoxyinosine and indole-3-carboxylate by exercise may have potentially contributed to antidepressant-like effects of exercise in CAF diet animals in the FST.

In conclusion, exercise produced a mild anxiolytic effect, regardless of diet, and increased PYY, a metabolic hormone previously shown to reduce anxiety-like behaviour. Furthermore, exercise mitigated CAF diet increases in immobility in the FST and in the metabolic hormones leptin and insulin, which have previously been implicated in depression-like behaviour. While a CAF diet robustly affected the differential expression of caecal metabolites, exercise attenuated a CAF diet-induced decrease in caecal abundance of anserine, deoxyinosine, and indole-3-carboxylate, metabolites previously implicated in cognitive function or depression-like behaviour. Although exercise exerted antidepressant-like effects in CAF diet-fed animals and induced subtle enhancements in spatial learning in standard chow-fed animals, we found that exercise needs to be accompanied by a healthy diet to increase AHN, possibly driven by alterations in the metabolic hormone GLP-1, thus highlighting the importance of exercise as well as a healthy diet for hippocampal health. Finally, these data provide additional insight into potential gut-mediated mechanisms underlying the effects of a CAF diet and exercise and shed light on potential targets for dietary supplementation to prevent a CAF diet from blunting beneficial effects of exercise on brain and behaviour.

## Author contributions

M.H.C. Nota, Y.M. Nolan, and O.F. O’Leary were responsible for conception, design and interpretation of the work. M.H.C. Nota. S. Nicolas, E.P. Harris and T. Foley acquired data. M.H.C. Nota and S. Dohm-Hansen analysed the data. M.H.C. Nota drafted the work, with assistance from S. Dohm-Hansen on caecal metabolomic methods and results sections. All authors have provided critical feedback for important intellectual content. given final approval of the version to be published and agree to be accountable for all aspects of the work.

## Acknowledgements

The authors thank Dr. Anna Golubeva and Dr. Gerard Moloney for their technical assistance.

---

**Supplementary Table 1:** Nutritional content of CAF foods and standard chow.

| Product | brand | per 100g |  |  |  |  |  |  |  |
| --- | --- | --- | --- | --- | --- | --- | --- | --- | --- |
|  |  | energy (kJ) | energy (kcal) | fat (g) | sat. fat (g) | carbs (g) | sugar (g) | fibre (g) | protein (g) |
| Frosted Shreddies | Nestle | 1567 | 370 | 1.5 | 0.2 | 76 | 25 | 9 | 9.2 |
| Choco Nut Pillows | Tesco | 1943 | 463 | 18.8 | 3.9 | 65.7 | 28.2 | 3.5 | 6 |
| Ripple Swiss Roll | Gateaux | 1442 | 342 | 7.5 | 2.7 | 65 | 42 | 0.8 | 3.6 |
| Jaffa Cakes | McVitie's | 1600 | 379 | 7.9 | 4 | 70.9 | 52.6 | 2.2 | 4.9 |
| Custard Cream Biscuits | Tesco | 2056 | 490 | 20.5 | 12.8 | 70.1 | 28.3 | 1.1 | 5.7 |
| KitKat (regular) | Nestle | 2101 | 502 | 24.4 | 13.7 | 62.7 | 51 | 2.1 | 6.7 |
| Milk Chocolate Digestives | Tesco | 2105 | 503 | 24.3 | 12.4 | 62.5 | 24.5 | 3.1 | 6.9 |
| Dry Roasted Peanuts | Tesco | 2581 | 623 | 51.7 | 9.5 | 10 | 2.9 | 6 | 26.5 |
| Walnuts | Alesto | 2938 | 712 | 69.1 | 6.8 | 3.7 | 3 | 6.8 | 15.5 |
| Double Cream Oreos | Mondelez | 2071 | 494 | 22 | 7.2 | 68 | 43 | 2.1 | 4.2 |
| Magic Stars | Milky way | 2330 | 559 | 35 | 22 | 54 | 54 | 0 | 6.4 |
| Cheese Twists | Tesco | 2120 | 5.7 | 26.4 | 14.5 | 52.1 | 2.5 | 2.8 | 13.8 |
| Lemon Cake | Tesco | 1744 | 416 | 18.2 | 2.4 | 59.6 | 40.4 | 1 | 2.9 |
| Chopped Pork and Ham | Dulano | 1117 | 269 | 22 | 8.8 | 2.6 | 1.4 | 0.5 | 15 |
| Ready Salted Crisps | Tesco | 2270 | 544 | 33.2 | 2.8 | 55.4 | 0.4 | 1.8 | 5 |
| Hula Hoops (cheese & onion) | KP Snacks | 2103 | 502 | 24 | 2.3 | 66 | 2.2 | 2.5 | 3.8 |
| Buttersalt Popcorn | Perri | 2252 | 542 | 36.4 | 4.7 | 40.8 | 0.1 | 9.9 | 7.9 |
| Mild White Cheddar Cheese | Tesco | 1619 | 390 | 32 | 20.8 | 0.1 | 0.1 | 0 | 25.5 |
| Frankfurters (pork) | Dulano | 1295 | 313 | 28 | 11 | 2 | 1 | 0.5 | 13 |
| Big Value Traditional Ham | Tesco | 416 | 99 | 2.4 | 0.9 | 0.5 | 0.5 | 0.5 | 18.5 |
| Jumbo Sausage Rolls | Lidl | 988 | 236 | 12.3 | 6 | 24.2 | 0.5 | 1.4 | 7.8 |
| Mini Rolls | Cadbury | 1925 | 460 | 23.5 | 11.3 | 56.3 | 42.8 | 2.3 | 4.8 |
| Tangy Cheese Doritos | Doritos | 2092 | 501 | 25.9 | 3.4 | 57.5 | 2.7 | 5.6 | 6.5 |
| Mini Battenbergs | Kipling | 1684 | 399 | 10 | 4 | 72.9 | 61.5 | 1.2 | 3.8 |
| M&Ms | Mars | 2020 | 481 | 19 | 12 | 70 | 68 | 0 | 5.2 |
| Jelly Babies | Sweet Corner | 1439 | 339 | 0.1 | 0.1 | 79.8 | 61.6 | 0.5 | 4.6 |
| Jelly Beans | Tesco | 1558 | 367 | 0.7 | 0.4 | 89.8 | 57.4 | 0.5 | 0.1 |
| Jelly Snakes | Natural Foods | 1384 | 326 | 0.2 | 0.2 | 77 | 61 | 0 | 3.1 |
| Standard chow | Envigo | 1300 | 310 | 6.2 | 0 | 44.2 | 0 | 3.5 | 18.6 |

**Supplementary Table 2:**
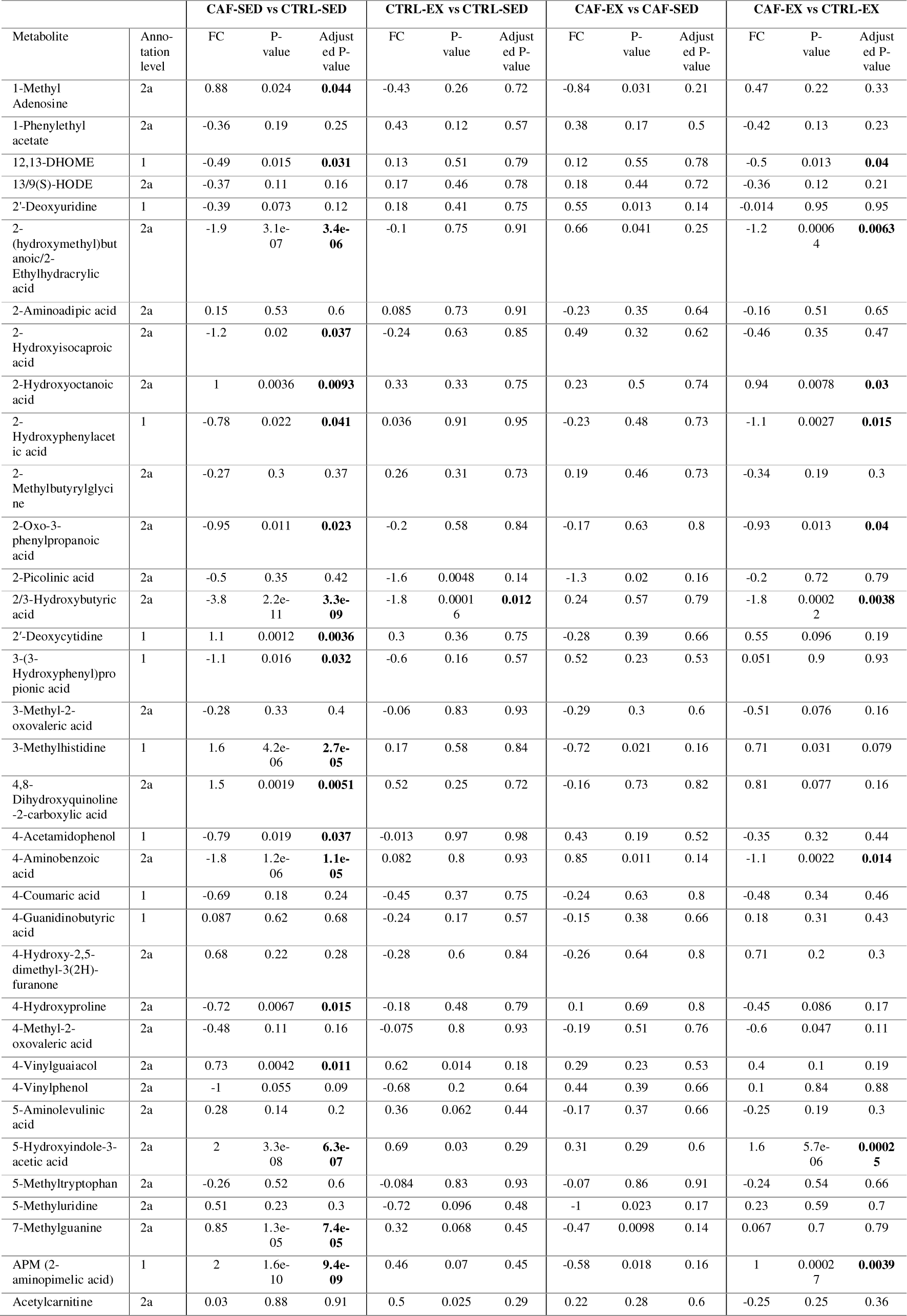
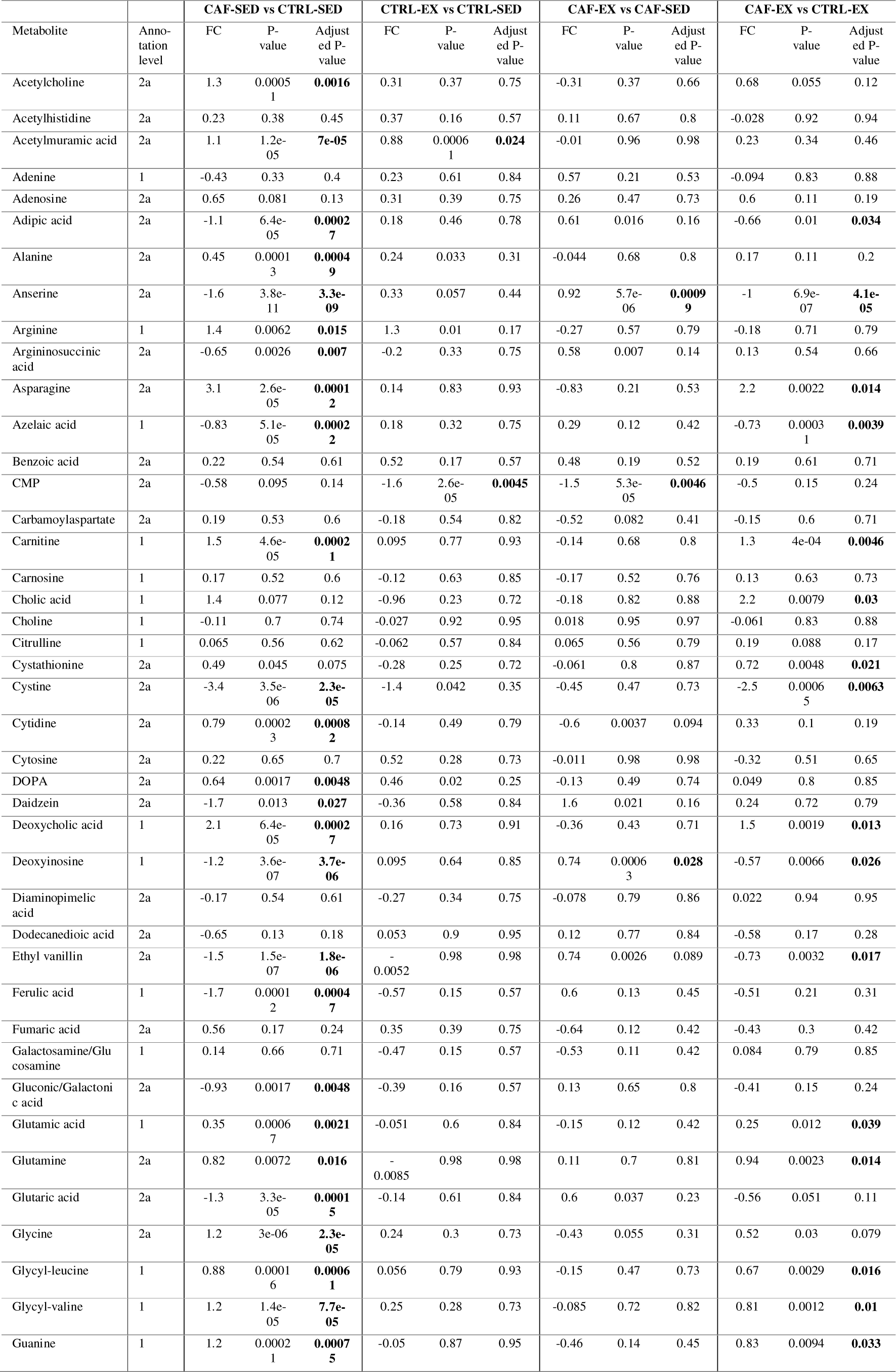
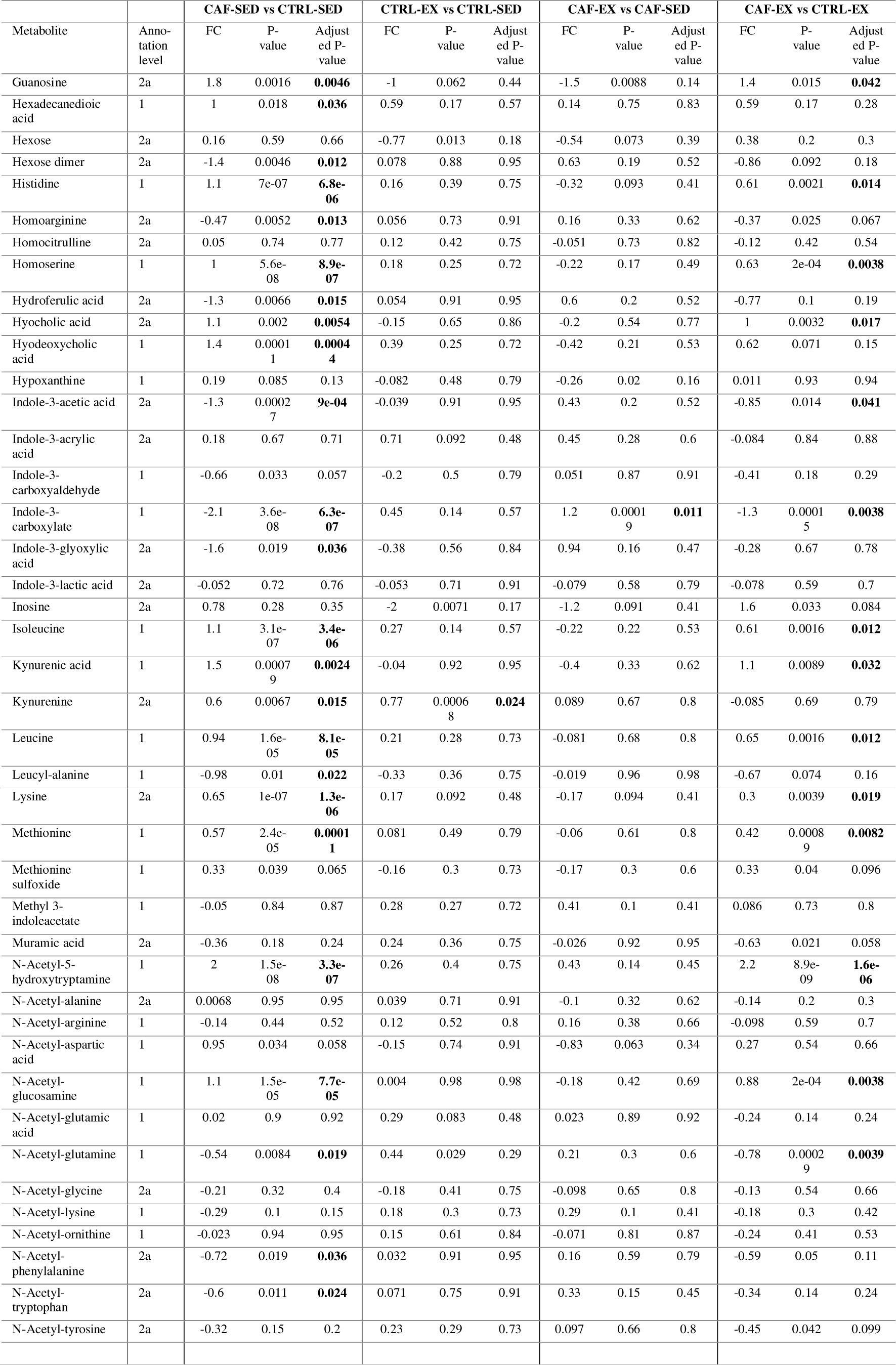
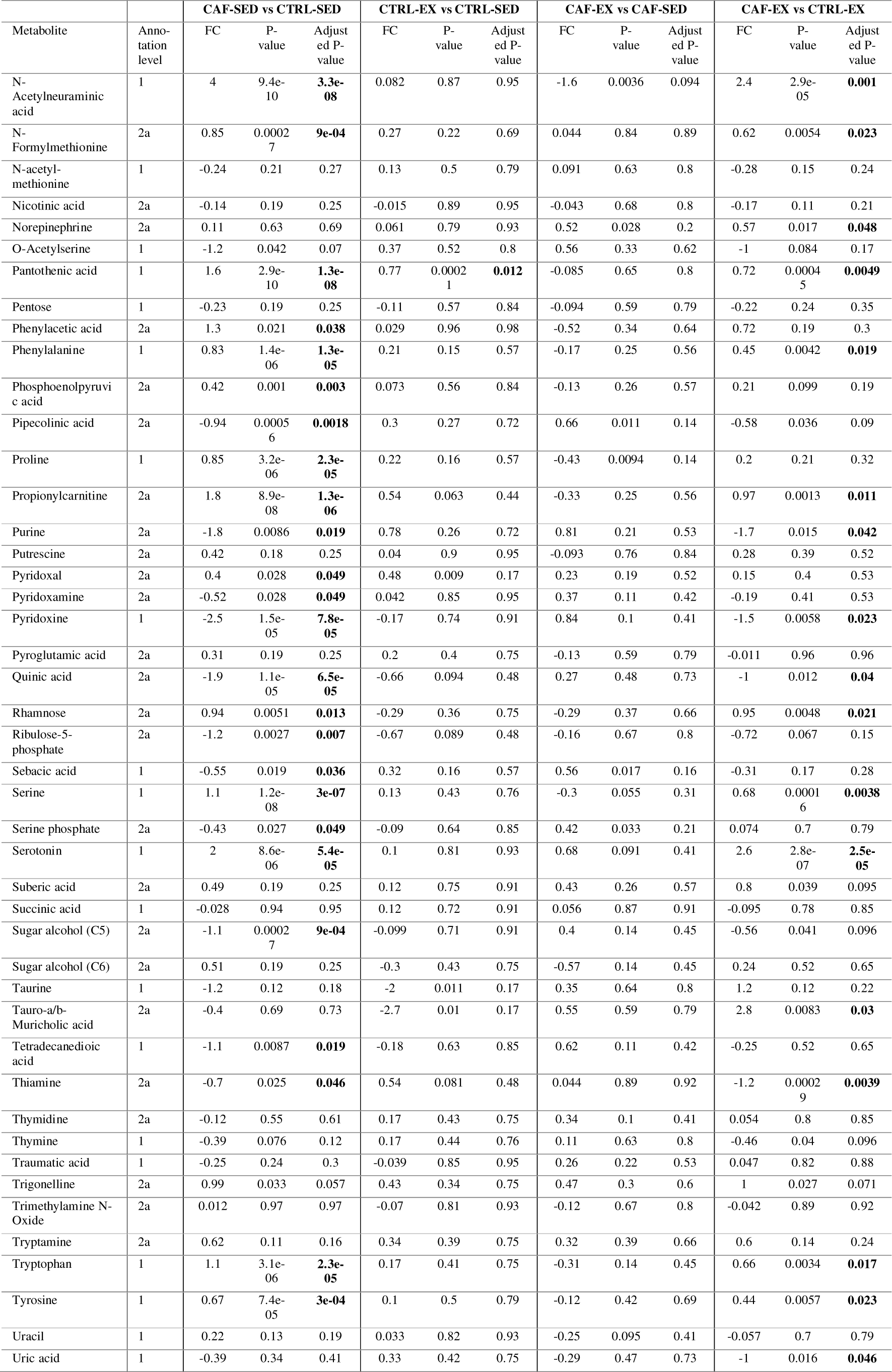
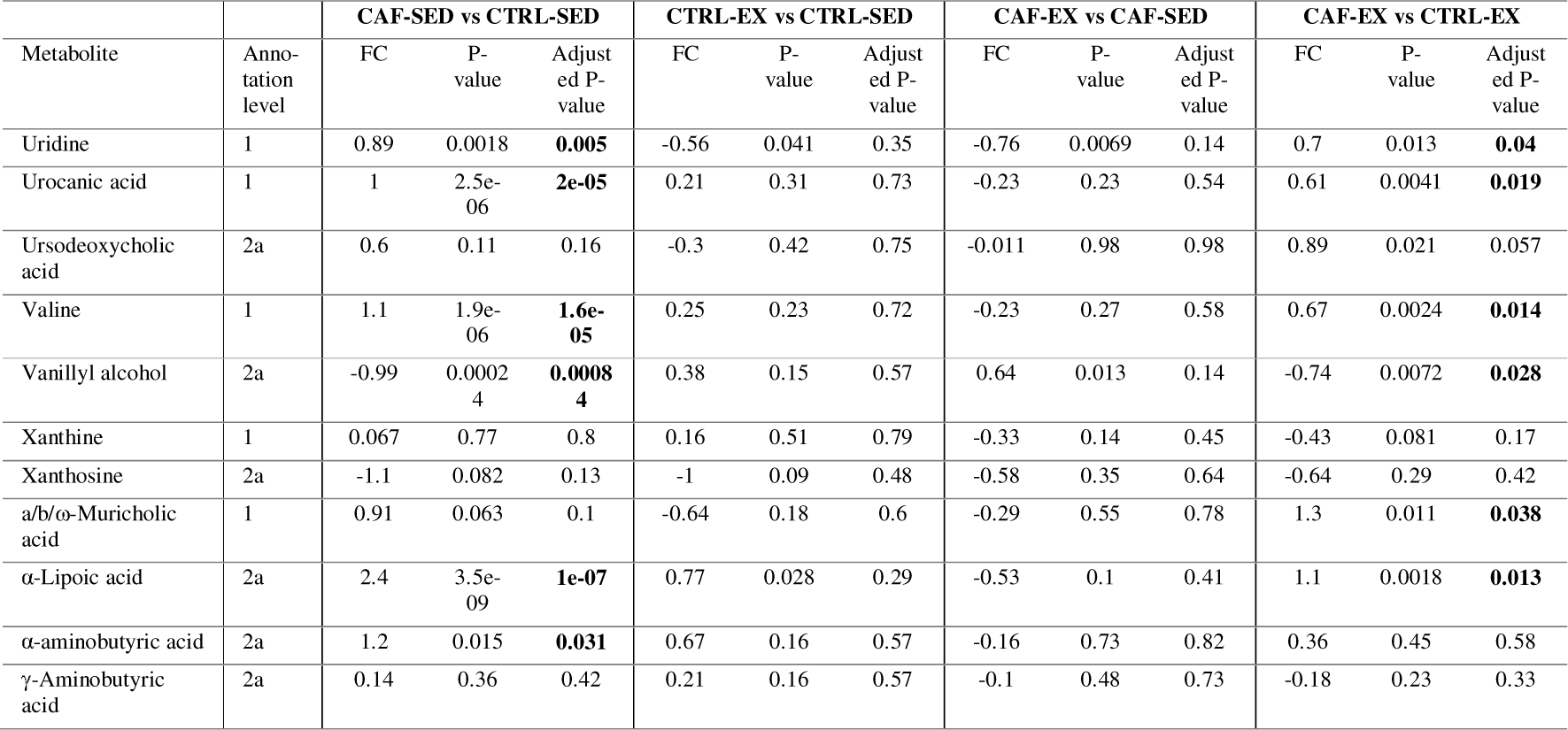
Differential expression analysis of quantified metabolites.

